# Detection of Ghost Introgression from Phylogenomic Data Requires a Full-Likelihood Approach

**DOI:** 10.1101/2023.04.29.538834

**Authors:** Xiao-Xu Pang, Da-Yong Zhang

**Affiliations:** State Key Laboratory of Earth Surface Processes and Resource Ecology and Ministry of Education Key Laboratory for Biodiversity Science and Ecological Engineering, College of Life Sciences, Beijing Normal University, Beijing 100875, China

## Abstract

In recent years, the study of hybridization and introgression has made significant progress, with ghost introgression - the transfer of genetic material from extinct or unsampled lineages to extant species - emerging as a key area for research. Accurately identifying ghost introgression, however, presents a challenge. To address this issue, we focused on simple cases involving three species with a known phylogenetic tree. Using mathematical analyses and simulations, we evaluated the performance of popular phylogenetic methods, including HyDe and PhyloNet/MPL, and the full-likelihood method, Bayesian Phylogenetics and Phylogeography (BPP), in detecting ghost introgression. Our findings suggest that heuristic approaches relying on site patterns or gene tree topologies struggle to differentiate ghost introgression from introgression between sampled non-sister species, frequently leading to incorrect identification of donor and recipient species. The full-likelihood method BPP using multilocus sequence alignments, by contrast, is capable of detecting ghost introgression in phylogenomic datasets. We analyzed a real-world phylogenomic dataset of 14 species of *Jaltomata* (Solanaceae) to showcase the potential of full-likelihood methods for accurate inference of introgression.

## Introduction

The growing availability of genomic data has led to an acceleration in research on hybridization and subsequent genetic exchange between species (i.e. introgression), which are widely recognized as significant factors in adaptation, speciation, and evolutionary innovation (Mallet et al. 2016; Taylor and Larson 2019; Edelman and Mallet 2021). In addition to ancient and contemporary gene flow between extant lineages extensively reported in the literature (Fontaine et al. 2015; Figueiró et al. 2017; Jones et al. 2018; Wu et al. 2018a; Zhang et al. 2019; Wang et al. 2020; Esquerré et al. 2021; Meleshko et al. 2021; Yang et al. 2021; Suvorov et al. 2022), the phenomenon of ‘ghost introgression’, which refers to gene flow from extinct or unsampled lineages to the sampled species, is gradually emerging as an important cutting-edge research topic (Ottenburghs 2020; Hibbins and Hahn 2022b; Tricou et al. 2022a, 2022b; Pang and Zhang 2023; Tiley et al. 2023). As the overwhelming majority of lineages have either gone extinct or have been unsampled due to technical limitations or irrelevance to specific research questions, evolutionary studies are inherently constrained to small subsets of species or populations. Ghost introgression, therefore, is a crucial factor to be reckoned with in such studies. Genomic data analyses have provided evidence of ghost introgression in plants (e.g., Ding et al. 2022; Li et al. 2022; Tiley et al. 2023) and animals (e.g., Ai et al. 2015; Kuhlwilm et al. 2019; Rocha et al. 2022), including humans (Green et al. 2010; Sankararaman et al. 2014).

Various phylogenetic methods have been developed over the past few years for detecting and characterizing introgression, ranging in complexity from heuristic methods based on summary statistics to full-likelihood techniques that utilize all the information present in multilocus sequence data (for latest reviews see Jiao et al. 2021; Hibbins and Hahn 2022b). Most of these methods use data from one sample per species, even when multiple samples are available (Hibbins and Hahn 2022b). These techniques were designed primarily to detect introgression between sampled living taxa, without considering the possibility of ghost introgression (Tricou et al. 2022a, 2022b). Evaluating these methods’ ability to detect ghost introgression will help researchers avoid misinterpretation of results and expand the tools available for detecting different types of gene exchange.

The most popular heuristic methods, such as the D-statistic (Green et al. 2010) and HyDe (Blischak et al. 2018), rely on summary statistics obtained from the site-pattern counts for a species quartet to test for the presence of gene flow between non-sister species. However, Tricou et al. (2022b) have shown that outgroup ghost introgression, defined as introgression from an unsampled lineage that diverged before the most basal species under investigation (illustrated in Fig. 1a), can significantly affect D-statistic values. This can lead to incorrect identification of both donor and recipient species involved in an introgression event. In an attempt to differentiate ghost introgression from other gene flow scenarios, Hibbins and Hahn (2022b) proposed using an additional species that is sister to the recipient but not implicated in any introgression as a reference. However, finding such a suitable reference species can be challenging, making this solution difficult to apply in practice. The HyDe test is based on a model of hybrid speciation and has been used as a well-justified approach for assessing introgression in general (Kong and Kubatko 2021). However, findings by Ji et al. (2023) suggest that the accuracy of HyDe may be compromised when inferring the donor and recipient of introgression events in outflow scenarios, where gene flow occurs from a sister species to the basal species (as illustrated in Fig. 1c). The behavior of HyDe under ghost introgression scenarios (Fig. 1a) has yet to be explored.

**Figure 1.**
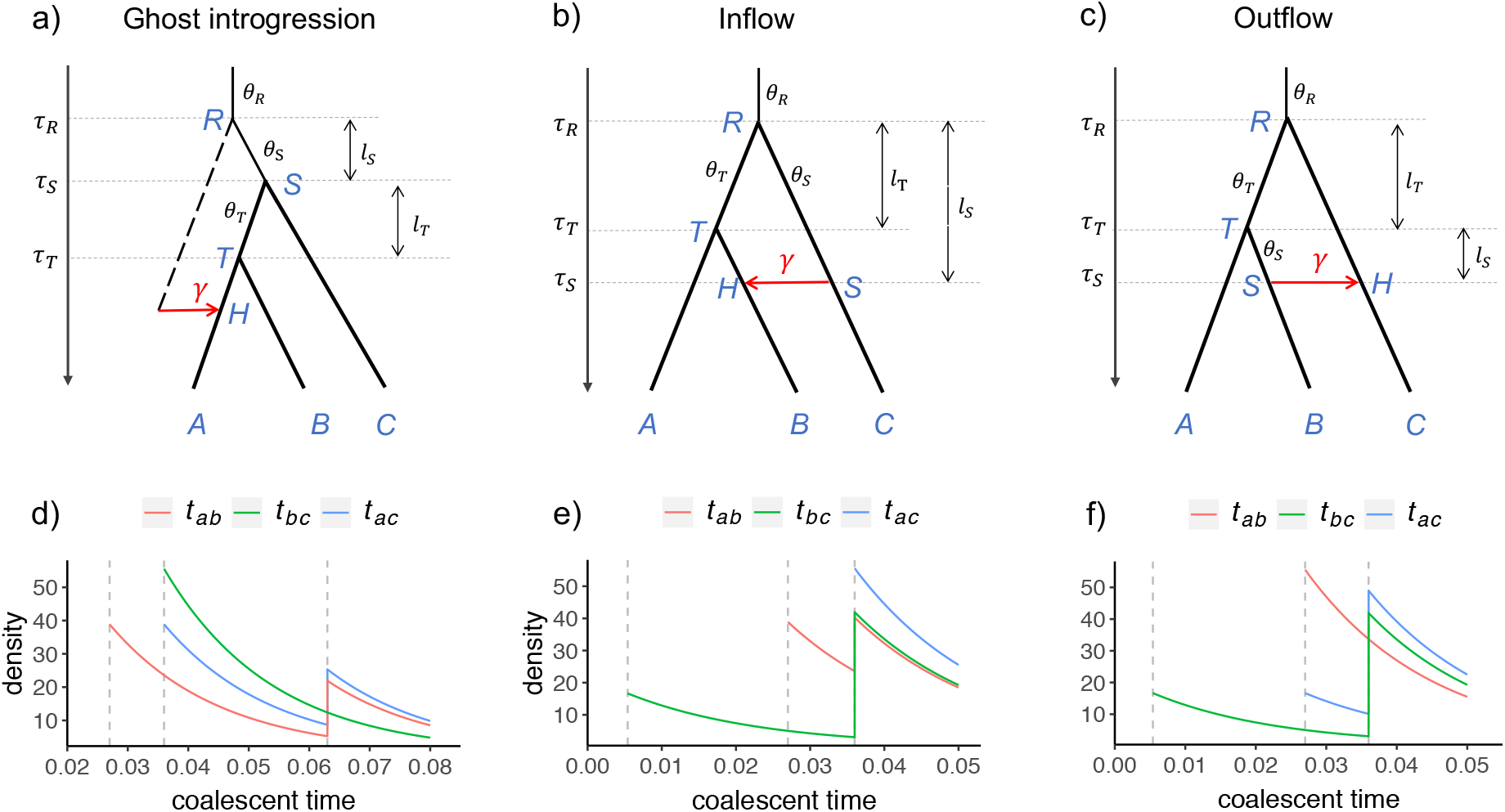
Three different introgression scenarios within a species tree *AB*|*C* and the corresponding distribution of coalescent times (*t*_*ab*_, *t*_*bc*_, *t*_*ac*_) for sequence pairs. a) Introgression from an outgroup ghost to sister species *A* (ghost introgression). b-c) Introgression between non-sister species in different directions *C*→*B* (inflow) and *B*→*C* (outflow). The backbone tree is represented by black thick branches, the introgression event is labelled by a red arrow, and the hybrid species *H* consists of a proportion *γ* of introgressed genes. *l*_*T*_ and *l*_*S*_ represent the branch length of the ancestral species *T* and *S*, respectively. d-f) The distribution of coalescent times (*t*_*ab*_, *t*_*bc*_, *t*_*ac*_) for sequence pairs in the introgression scenarios shown above in Figures 1a-c. The parameters in the ghost introgression scenario are {*θ*_*S*_ = *θ*_*T*_ = *θ*_*R*_ = *θ* = 0.036, *τ*_*T*_ = 0.75*θ, τ*_*S*_ = *θ, τ*_*R*_ = 1.75*θ, γ* = 0.3}. The parameters in the inflow and outflow cases are {*θ*_*S*_ = *θ*_*T*_ = *θ*_*R*_ = *θ* = 0.036, *τ*_*S*_ = 0.15*θ, τ*_*T*_ = 0.75*θ, τ*_*R*_ = *θ, γ* = 0.3}.

Another class of approaches is designed to detect introgression across an entire phylogeny by estimating a phylogenetic network on the full set of taxa under investigation. Heuristic methods for constructing these networks use information from gene tree topologies (or subtree topologies for subsets of species trios or quartets) and then analyze them using a (pseudo)likelihood or distance framework. Examples of such methods include InferNetwork_MPL and InferNetwork_ML in the PhyloNet program (Yu et al. 2014; Yu and Nakhleh 2015), SNaQ in the PhyloNetworks package (Solís-Lemus and Ané 2016) and NANUQ in the MSCquartets package (Allman et al. 2019). However, these heuristic methods have a significant drawback: networks may not always be identifiable. This means that the information derived solely from gene tree topologies may be insufficient to differentiate between networks representing different biological hypotheses about speciation and introgression (Pardi and Scornavacca 2015; Zhu and Degnan 2017; Yang and Flouri 2022).

Full-likelihood phylogenetic network methods refer to those that use the joint distribution of gene trees and coalescent times and operate on multilocus sequence data directly (Jiao et al. 2021). Due to their accommodation of the information in gene-tree branch length (coalescent times), full-likelihood methods significantly improve statistical power in identifying networks, but at the expense of a much-increased computational burden (Hey et al. 2018; Wen and Nakhleh 2018; Zhang et al. 2018a; Flouri et al. 2020). The Bayesian Phylogenetics and Phylogeography (BPP) program (Flouri et al. 2020) is notable in allowing users to compare two putative networks using Bayes factor, avoiding cross-model search in large network space. Thus it is much more computationally feasible.

Here we examine the effectiveness of various phylogenetic techniques in detecting ghost introgression through a study of three introgression scenarios that can lead to the same significant D-statistic, as first made explicit by Tricou et al. (2022b). These scenarios include outgroup ghost introgression (Fig. 1a) and ingroup introgression between non-sister species, with either *C* → *B* (inflow, Fig. 1b) or *B* → *C* (outflow, Fig. 1c) directions within the species tree *AB*|*C*. Our analysis is limited to cases where only one sequence is sampled from each species, as the addition of more samples may provide little new information regarding introgression (Hibbins and Hahn 2022b). We begin by mathematically deriving probabilities of gene tree topologies, frequencies of biallelic site patterns, and distributions of coalescent times for pairs of sequences from different species. This allows us to analyze how well these pieces of information can distinguish outgroup ghost introgression from other introgression scenarios. We then conduct simulations to examine the behaviors of site pattern-based HyDe, gene tree-based InferNetwork_MPL in PhyloNet (or simply PhyloNet/MPL), and the full-likelihood method BPP under the aforementioned introgression scenarios. As a real-world example, we analyze a phylotranscriptomic dataset of 14 *Jaltomata* (Solanaceae) species (Wu et al. 2018b; Tiley et al. 2023) to demonstrate the potential pitfalls of using heuristic methods as well as the feasibility of using full-likelihood methods to detect ghost introgression in mid-sized groups of lineages of interest. By integrating theoretical analysis, simulation evaluation, and practical examples, our study provides valuable insights for biologists investigating ghost introgression, highlighting which methods are most suitable for this very purpose.

## Theory

We consider three gene flow scenarios, likely associated with a significant D-statistic (Tricou et al. 2022b), within the species tree *AB*|*C* (along with a distant outgroup *O*) under the model of multispecies coalescent with introgression (MSci; Yu et al. 2012). These scenarios include outgroup ghost introgression involving gene flow from an outgroup ghost lineage to species *A* (Fig. 1a, hereafter referred to as ghost introgression), as well as ingroup introgression between non-sister species *B* and *C* in different directions (inflow and outflow, depicted in Figs. 1b-c). Both species divergence times (*τ*_*R*_, *τ*_*T*_, *τ*_*S*_) and population sizes (*θ*_*R*_, *θ*_*T*_, *θ*_*S*_) in the models are measured by the expected number of mutations per site; i.e., one time unit is the expected time to accumulate one mutation per site. Specifically, the divergence time is defined as *τ* = *Tμ*, where *T* is the divergence time in generations and *μ* is the mutation rate per site per generation. The population size refers to *θ* = 4*Nμ*, where *N* is the effective population size of the species. The data analyzed consist of multiple loci, with three sequences from each species at each locus (*a, b, c*). The possible gene-tree topologies at each locus are *G*_1_ = *ab*|*c, G*_2_ = *a*|*bc* and *G*_3_ = *ac*|*b*.

Note that the three scenarios of ghost introgression, inflow, and outflow share the same speciation history of *AB*|*C* and the introgressed history of *A*|*BC*, despite having distinct recipients of gene flow - *A, B*, and *C*, respectively. As a result, the three introgression scenarios can produce similar features in gene genealogies and in sequence data. Our main focus is to examine to what extent the three introgression scenarios can be distinguished with different types of information. Specifically, we derive analytical results in network identification and the estimation of introgression probability *γ* for three methods: 1) PhyloNet/MPL, which is based on probabilities of gene-tree topologies, 2) HyDe, which relies on site-pattern frequencies, and 3) full-likelihood methods, which use the joint distribution of gene tree topologies and coalescent times.

### Identifiability based on Probabilities of Gene-tree Topologies (Analysis of PhyloNet/MPL)

In the three MSci models shown in Figure 1, we define *C*_*T*_ = 2*l*_*T*_/*θ*_*T*_ and *C*_*S*_ = 2*l*_*S*_/*θ*_*S*_ as the lengths, in coalescent units, of the internal branch *T* in the species history and of the internal branch *S* in the introgression history, respectively. Following the speciation history *AB*|*C* with a probability of 1 − *γ*, if sequences *a* and *b* coalesce in species *T*, the gene tree would be *G*_1_ = *ab*|*c*; following the introgressed history *A*|*BC* with a probability of *γ*, if sequences *b* and *c* coalesce in species *S*, the gene tree would be *G*_2_ = *a*|*bc*. If neither of these events occurs, indicating the occurrence of incomplete lineage sorting (ILS) within either the speciation or introgressed history, then three gene trees will occur with equal probability. The probabilities of the three gene-tree topologies can be represented using the following equations, which have been derived in Jiao et al. (2020) and Pang and Zhang (2023):

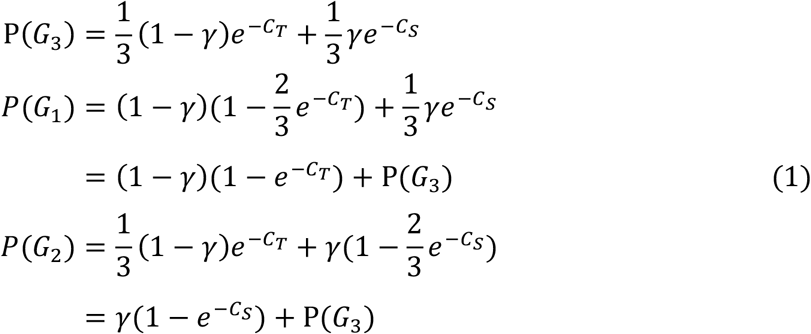

The sum of three gene-tree topology probabilities equals 1, leaving only two free quantities: 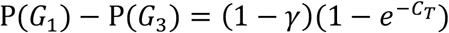 and 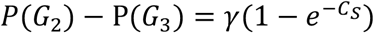. It is worth noting that the expressions for gene-tree topology probabilities can be applied to all three introgression scenarios. This means that gene-tree topologies can have identical probabilities across the three introgression scenarios, provided that the values of 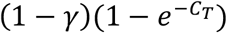 and 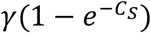 remain invariant across scenarios. Consequently, the PhyloNet/MPL method that uses information only from gene tree topologies cannot differentiate between the scenarios of ghost introgression, inflow, and outflow. Moreover, even if multiple sequences per species are utilized, the issue with PhyloNet/MPL remains unresolved. The reason why PhyloNet/MPL cannot improve network identification is because it only considers sequence permutations within three species, selecting one sequence from each species. Thus, this approach results in only three gene tree topologies without providing more information to enhance network inference.

Furthermore, we examine the estimation of introgression probability *γ*. With only two independent equations, there are infinite solutions for the three unknown parameters {*γ, C*_*T*_, *C*_*S*_}. Specifically, given the observed gene-tree topology probabilities *P*(*G*_1_), *P*(*G*_2_) and *P*(*G*_3_), any value of *γ* within the range of *P*(*G*_2_) − *P*(*G*_3_) < *γ* < 1 − (*P*(*G*_1_) − *P*(*G*_3_)) can find corresponding solutions for *C*_*T*_ and *C*_*S*_ to satisfy the two equations. By substituting the expressions for *P*(*G*_*i*_) {*i* = 1, 2, 3} from equation 1, it can be shown that the PhyloNet/MPL estimates of *γ*, denoted as *γ*_*phy*_, will fall within this range:

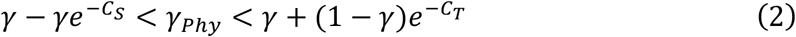

Obviously, the upper and lower bounds for *γ*_*phy*_ depend respectively on the value of *C*_*T*_, reflecting the level of ILS within the speciation history, and the value of *C*_*S*_, reflecting the level of ILS within the introgressed history (Figs. 2a-b). The larger values of *C*_*T*_ and *C*_*S*_, and hence the weaker ILS in both histories, the more precise estimation of *γ* will be achieved. However, when both *C*_*T*_ and *C*_*T*_ approach zero, the estimated *γ* will become highly variable within the range of 0 to 1 (Fig. 2a-b).

**Figure 2.**
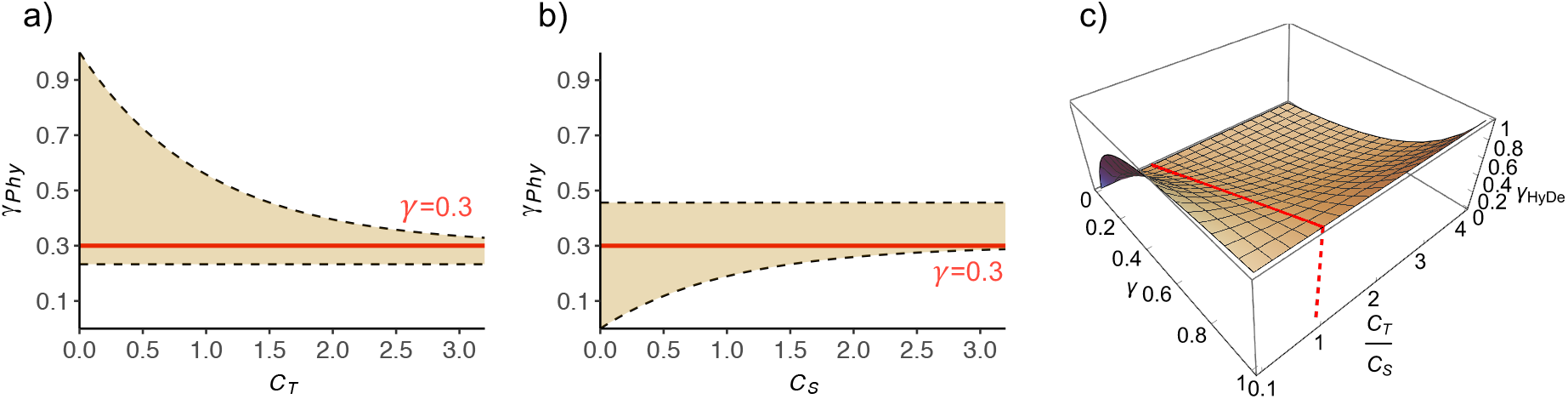
Estimation of introgression proportion *γ* in PhyloNet/MPL and HyDe. In reference to the introgression scenarios illustrated in Figure 1, *C*_*T*_ = 2*l*_*T*_/*θ*_*T*_ and *C*_*S*_ = 2*l*_*S*_/*θ*_*S*_ denote the branch lengths in coalescent units of ancestral species *S* and *T*, respectively. a-b) In PhyloNet/MPL, when *γ* = 0.3 labelled by the red line, the estimates *γ*_*phy*_ would disperse over the range demarcated by the yellow shade, with the upper and lower bounds labelled by the dashed lines, correspondingly. a) The range for *γ*_*phy*_ varies with the value of *C*_*S*_ after fixing *C*_*T*_ = 1.5. b) The range for *γ*_*phy*_ varies with the value of *C*_*S*_ after fixing *C*_*T*_ = 1.5. Obviously, the upper and lower limits for estimated *γ*_*phy*_ depend respectively on the value of *C*_*T*_ and *C*_*S*_. c) In HyDe, under the assumption of constant population size, the estimated *γ*_HyDe_ is related to the ratio *C*_*T*_/*C*_*S*_. When *C*_*T*_ = *C*_*S*_, HyDe can accurately estimate the value of *γ*, as demonstrated by the red line. Otherwise, *γ* may be underestimated or overestimated, depending on the relative values of *C*_*T*_ and *C*_*S*_.

### Identifiability based on Site Patterns (Analysis of HyDe)

Assuming an infinite-site mutation model (i.e., no multiple hits), three biallelic site patterns, ABBA, AABB and ABAB (with A representing the ancestral state from the outgroup that is listed first and B the derived state), are generated through mutations on the internal branches of gene trees *G*_1_, *G*_2_, and *G*_3_, respectively. Thus, the expected number of the three parsimony-information sites can be indicated by the sum of the lengths of internal branches in the relevant gene trees (Mendes and Hahn 2018; Hibbins and Hahn 2022a). Specifically, we weight the internal branch lengths of the relevant gene trees by their probabilities and sum them across coalescent histories (more details are provided in Supplementary Note 1). Here, we derive the frequency of each site pattern per nucleotide site under the assumption of a constant population size with a common *θ*:

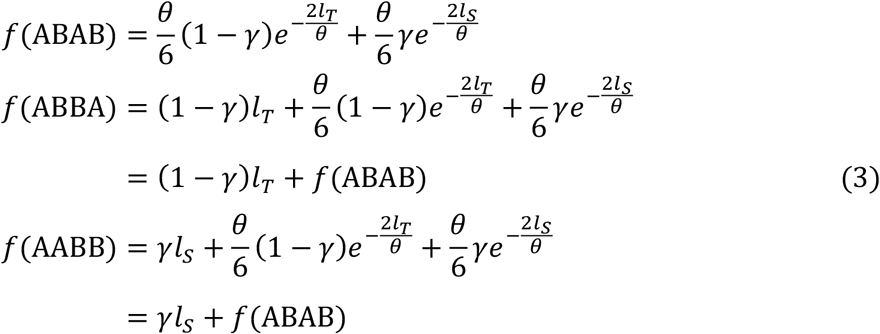

Hibbins and Hahn (2022a) have derived similar equations by considering branch lengths in coalescent units. Notably, these expressions are universally applicable to all three introgression scenarios, where the parameters *l*_*T*_ and *l*_*S*_ denote the lengths of the internal branches within the speciation and the introgressed histories, respectively, in each scenario. In all three scenarios, the site patterns are ordered in the same way, i.e., *f*(ABAB) <*min*{*f*(ABBA), *f*(AABB)}.

HyDe executes the statistical test for introgression detection, with the null hypothesis being the MSC model where the two smallest site patterns are equal (Kubatko and Chifman 2019). Thus, the effectiveness of HyDe in detecting introgression is related to the value of *min*{*f*(ABBA) − *f*(ABAB), *f*(AABB) − *f*(ABAB)}, i.e., *min*{(1 − *γ*)*l*_*T*_, *γl*_*S*_}, with a larger value indicating greater ease of detection. HyDe interprets a significant outcome as resulting from a hybrid speciation event, in which the two parent species are expected to be more distantly related than either of them to the hybrid. Thus, the two species that share the derived state in the smallest number of site patterns will be identified as putative parents, and the remaining species as a hybrid (Kubatko and Chifman 2019). In the three distinct introgression scenarios in Figure 1a-c, the site pattern ABAB is always least numerous; as a result, HyDe will invariably point to an inflow scenario with species *B* being the hybrid. Finally, HyDe estimates *γ* using the function (Blischak et al. 2018):

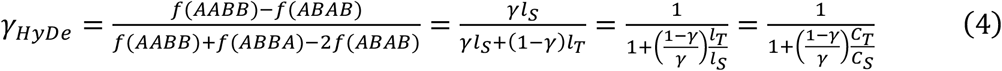

The estimation of *γ* is tied to *l*_*T*_/*l*_*S*_ or *C*_*T*_/*C*_*S*_. Accurate estimation of *γ* can only be achieved when *l*_*T*_ = *l*_*S*_ (or *C*_*T*_ = *C*_*S*_). Otherwise, the value of *γ* will either be underestimated or overestimated, depending on whether the ratio of *l*_*T*_/*l*_*S*_ (or *C*_*T*_/*C*_*S*_) is larger or smaller than 1 (Fig. 2c).

### Network Identifiability of Full-Likelihood Methods

Full-likelihood methods use the joint probability distribution of gene tree topologies and coalescent times (i.e., gene-tree branch lengths). For ease of analysis, we focus on the distributions of coalescent times between any two sequences, namely, *t*_*ab*_, *t*_*bc*_ and *t*_*ac*_ (Fig. 1d-f). In a population of size *θ*, the coalescent time of two sequences follows an exponential distribution with a mean of *θ*/2. The probability densities of coalescent times for the three introgression scenarios are derived in Supplementary Note 2 and plotted for specific parameters in Figures 1d-f.

There are two unique characteristics of coalescent times for each scenario. Firstly, the order of the minimum coalescence times for the three sequence pairs differs across the three cases. Specifically, in the ghost introgression scenario, the minimum coalescence times of sequence pairs align with the original species tree, where min(*t*_*ab*_) < min(*t*_*ac*_) = min(*t*_*bc*_) (Fig. 1d). In the inflow case, introgression reduces the minimum coalescent time of the sequence pair *bc* from the root time to the introgression time, while the minimum coalescent times for the other two sequence pairs remain unaffected. As a result, the order becomes min(*t*_*bc*_) < min(*t*_*ab*_) < min(*t*_*ac*_) (Fig. 1e). In the outflow case, introgression reduces the minimum coalescent times for sequence pairs *ac* and *bc* from the root time to the speciation time of the sister species and to the more recent introgression time, respectively, resulting in the order of min(*t*_*bc*_) < min(*t*_*ab*_) = min(*t*_*ac*_) (Fig. 1f).

Secondly, the distribution form of coalescence times for sequence pairs varies among the three scenarios of ghost introgression, inflow, and outflow, each of which involves a specific recipient: *A, B*, and *C*, respectively. The coalescent time of the sequences of the recipient and another species will have a mixture distribution that is discontinuous at certain time points because the sequence of the recipient lineage has two coalescence paths. However, introgression does not affect the coalescent process for a pair of sequences from the two species not acting as the recipient, and the coalescent time follows a smooth shifted exponential distribution when constant population size is assumed. Thus, using one sequence per species, full-likelihood methods are capable of distinguishing among the three scenarios.

## Simulations

### Data Simulation for Evaluation of Introgression Detecting Methods

We simulated three scenarios - ghost introgression, inflow, and outflow - with varying levels of ILS within the speciation history (*C*_2_), as well as varying levels (*γ*) and times (*C*_1_) of introgression, as shown in Figure 3. For each case, we set the linking of the divergence of outgroup species *O* to *R* to five coalescent units, and we simulated 100 replicates. We employed *ms* simulator (Hudson 2002) to generate 1,000 gene trees with one sequence per species. These gene trees were then used to simulate DNA sequences of 1,000 base pairs by Seq-Gen (Rambaut and Grass 1997) under a rate of population mutation of *θ* =0.036 (as in Yu et al. 2014; Kong and Kubatko 2021) and the JC model. The concatenated sequences were used for HyDe. In particular, we found that HyDe has a high false positive rate due to the non-independence of sites within a locus sharing the same underlying genealogy (Supplementary Figure 1). To address this issue, we used the block-jackknife approach to estimate sample variance and then to calculate the significance value. We chose the 1% significant level to conduct tests as it resulted in a low false positive rate (Supplementary Figure 1). We rerooted the simulated gene trees using the outgroup *O*, which was then removed. The rooted gene trees were used as inputs for the maximum pseudo-likelihood method InferNetwork_MPL in PhyloNet v3.8.2 (PhyloNet/MPL). Furthermore, we conducted additional simulations with four sequences per species to explore the impact of multiple sequences per taxon on PhyloNet/MPL.

**Figure 3.**
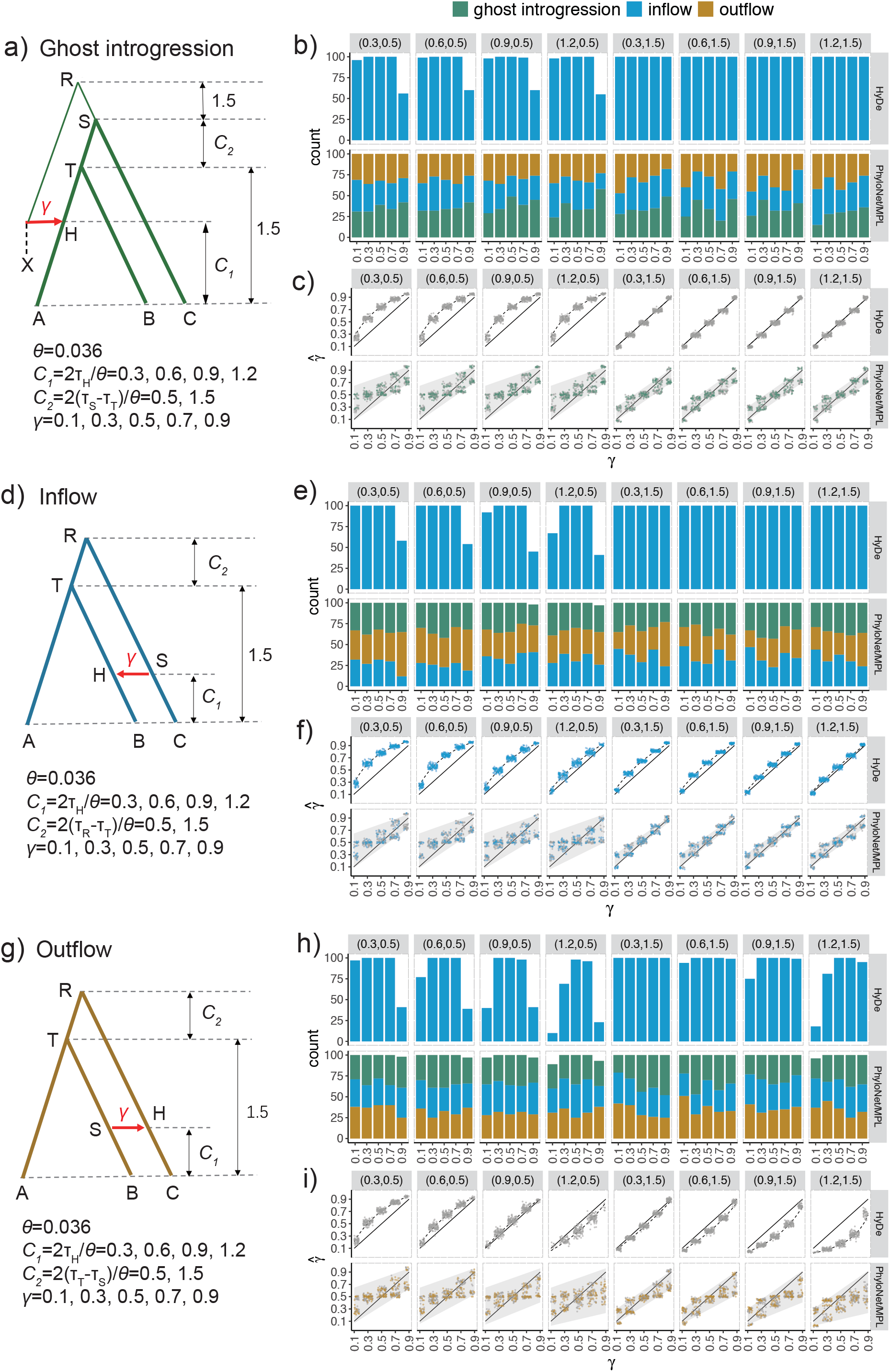
Results of HyDe and PhyloNet/MPL for the scenarios of ghost introgression, inflow, and outflow. a) Ghost introgression scenarios with corresponding parameter settings. b-c) Plots of results for the network inference and *γ* estimation for ghost introgression simulations. The strips located on the top and right of each plot represent branch lengths in coalescent units (*C*_1_, *C*_2_) and the methods employed, respectively. The x-axis indicates the value of *γ*. b) Estimation of network topologies. The numbers of topologies of three network models (i.e., ghost introgression, inflow and outflow) inferred by HyDe and PhyloNet/MPL (using the command InferNetwork_MPL) among 100 replicates are represented by colored bars. The remaining replicates that are not shown correspond to statistical insignificant results in HyDe or other networks in PhyloNet/MPL. c) Estimation of introgression probability *γ* by HyDe and PhyloNet/MPL. Colored and grey points represent the estimates of *γ* in the true and other two false networks, respectively. These points are horizontally jittered to avoid clutter. The solid line shows the true values, while the dashed line displays the expected estimates by HyDe based on equation 4 in the main text, and grey shade area represents the expected range of estimation by PhyloNet/MPL according to equation 2 in the main text. d-f) Inflow scenarios and corresponding simulation results. g-i) Outflow scenarios and corresponding simulation results.

We used the full-likelihood implementation of the MSci model in BPP v4.6.2 (Flouri et al. 2020) to analyze the sequence data (excluding outgroup *O*). There are other full-likelihood methods available in literature, including MCMC_SEQ in PhyloNet (Wen and Nakhleh 2018) and Species Network in *BEAST (Zhang et al. 2018a), but they are too computationally demanding to use in the present simulation study. In BPP, we conducted comparisons of ghost introgression, inflow and outflow scenarios by calculating the marginal likelihood. Thermodynamic integration combined with Gaussian quadrature was used to calculate the marginal likelihood under each scenario, with 16 quadrature points (Lartillot and Philippe 2006; Rannala and Yang 2017). For each of 16 MCMC runs, we used 50,000 MCMC iterations as burnin and then took 10^6^ samples, sampling every 2 iterations. Then, we conducted A00 analysis (Yang 2015) to estimate the parameters under the true MSci model with the same MCMC iteration settings. Each run took ∼6 hrs using single thread.

We further evaluated the performance of PhyloNet/MPL in detecting ancient ghost introgression events. We considered ancient introgression scenarios where lineages diverge subsequent to an introgression event (Fig. 5). For each condition, we simulated 20 replicates and generated 1,000 gene trees for each replicate with one sequence per species. The gene trees were rerooted by the outgroup *O*. These rooted gene trees without the outgroup were used as inputs for PhyloNet/MPL.

### Results for the Heuristic Methods: HyDe and PhyloNet/MPL

The results of HyDe and PhyloNet/MPL under different combinations of *C*_1_, *C*_2_, and *γ* in ghost introgression scenarios are summarized in Figures 3a-c. The sensitivity of HyDe for detecting introgression is not influenced by the time of introgression (*C*_1_), but rather by the degree of ILS in the speciation history (*C*_2_) and the level of introgression (*γ*), as shown in Figure 3b. A higher ILS in the speciation history (smaller *C*_2_) diminishes the power of HyDe for introgression detection. Moreover, as theoretically expected, HyDe wrongly attributed ghost introgression to inflow. The recipient of gene flow, *A*, is mistakenly identified as the donor, while its sister, *B*, is identified as a hybrid lineage. The performance of PhyloNet/MPL is also poor, as it has a tendency to randomly infer three scenarios of ghost introgression, inflow, and outflow, regardless of the parameter combinations (Fig. 3b). This result is in line with our theoretical analysis, which suggests that the probabilities of gene tree topologies are not capable of distinguishing between the three introgression scenarios. Regarding the introgression probability *γ*, the estimates in HyDe conform perfectly to the theoretical prediction derived from equation 4, even though the assumption of an infinite site model is not strictly satisfied in our simulation data (Fig. 3c). The *γ* estimates obtained from PhyloNet/MPL fall within the range predicted by equation 2. More specifically, the accuracy of *γ* estimation in HyDe and PhyloNet is linked to the level of ILS within the speciation history (*C*_2_), as well as the amount of ILS within the introgression history that depends on the distance of the outgroup ghost (i.e., the length of *RS* fixed at 1.5 in Fig. 3a). When ILS is strong in the speciation history with *C*_2_ = 0.5, HyDe overestimates *γ* because *C*_2_ < *RS*. The estimates from PhyloNet/MPL exhibit multimodality, forming two clusters with a large difference between them. In the cases with *C*_2_ = 1.5, HyDe accurately estimates *γ*, as the levels of ILS within the two histories are equal, despite an incorrect inference of the introgression scenario. In PhyloNet/MPL, the difference between the two clusters of estimates decreases, and the estimates become closer to the true value. Notably, the dispersion of *γ* estimation in PhyloNet/MPL is not caused by random error resulting from insufficient data, but rather by systematic bias arising from parameter unidentifiability.

The results of the inflow scenario are summarized in Figures 3d-f. HyDe performs well in this case because it accurately identifies the hybrid lineage *B* in almost all replicates except when ILS is prevalent at *C*_2_ = 0.5 and *γ* = 0.1, 0.9, in agreement with the results of Kong and Kubatko (2021). By contrast, PhyloNet/MPL performs poorly and cannot distinguish the three scenarios (Fig. 3e). HyDe overestimates the introgression probability *γ* (Fig. 3f), as the level of ILS within the speciation history is always higher than that within the introgressed history (*RT* < *RS*) under a constant population size. The extent of overestimation is greater when ILS is stronger in the speciation history and the introgression event happened more recently, e.g., (*C*_1_, *C*_2_) = (0.3, 0.5). The precision of *γ* estimates in PhyloNet/MPL is mainly influenced by the extent of ILS within the speciation history (*C*_2_). When ILS is prevalent such as when *C*_2_ = 0.5, the estimates are dispersed over a broad range, suffering from high variance. In contrast, the estimates of *γ* become more precise and stable when *C*_2_ = 1.5.

Figures 3g-i present the results of the outflow cases. HyDe is slightly less powerful in detecting introgression (Fig. 3h) in these cases than in inflow cases (Fig. 3e). The power is less than 50% in certain cases with high ILS within the speciation history (*C*_2_ = 0.5) and extreme values of introgression probability (*γ* = 0.1, 0.9). More significantly, HyDe reverses the direction of introgression and misidentifies species *B* as hybrid, as also found in the simulation of Ji et al. (2023). *γ* is overestimated by HyDe in some cases, such as when (*C*_1_, *C*_2_) = (0.3, 0.5), and underestimated in other cases, such as when (*C*_1_, *C*_2_) = (1.2, 1.5), depending on the relative levels of ILS within the speciation (*C*_2_) and introgressed histories (the length of the branch *TS*) (Fig. 3i). When the levels of ILS are close, such as at (*C*_1_, *C*_2_) = (0.9, 0.5), even though the direction of gene flow between non-sister species is incorrectly inferred, the extent of introgression *γ* is estimated with high accuracy. PhyloNet/MPL easily confuses outflow with ghost introgression or inflow (Fig. 3h), and exhibits a great degree of variability for estimates of *γ* at a high level of ILS (Fig. 3i).

We also investigated the performance of PhyloNet/MPL when multiple sequences were used per species, with results summarized in Figures S2-4. The performance of PhyloNet/MPL in such cases was similar to the single-sequence cases. In other words, even when multiple sequences per species are available, PhyloNet/MPL is still unable to differentiate between the three introgression scenarios. Thus we conclude that PhyloNet/MPL generally cannot identify ghost introgression.

### Results for the Full-likelihood Method: BPP

Due to considerable computational requirements, we limited our analysis to the parameter combination (*C*_1_, *C*_2_, *γ*) = (0.3, 0.5, 0.3) for three scenarios of ghost introgression, inflow and outflow. We compared three network models by Bayes factor through BPP for each simulation data, and the results are presented in Figure 4. For each scenario, it was observed that the true introgression model exhibited substantial superiority over other models, with the logarithm of Bayes factors exceeding 22 across all replicates (Fig. 4a). This means that the ghost introgression and ingroup introgressions (i.e., inflow and outflow) confused by heuristic methods can be successfully distinguished by full-likelihood analysis making full use of both topological and branch-length information of gene trees. Moreover, in comparison with heuristic approaches, full-likelihood methods can identify the parameters of species divergence times *τ*_*S*_, ancestral population sizes *θ*_*S*_, and introgression probability *γ* (Figs. S5 and 4b). In all three scenarios, the introgression probability *γ* and species divergence times (*τ*_*S*_, *τ*_*T*_, *τ*_*R*_) are well-estimated with narrow confidence intervals (CI). However, it should be noted that the introgression time *τ*_*H*_ cannot be inferred here, but it would become identifiable when using multiple sequences per species. The precision of the estimated population size *θ*_*S*_ varies across different ancestral populations. In all scenarios, the population sizes of species *T* and *S* (*θ*_*T*_ and *θ*_*S*_) are estimated poorly when compared to those of species *R* (*θ*_*R*_), which can be explained by the fact that more coalescence events occur in species *R* than in species *S* and *T*. More precise results are expected with an increasing number of loci (Huang et al. 2020).

**Figure 4.**
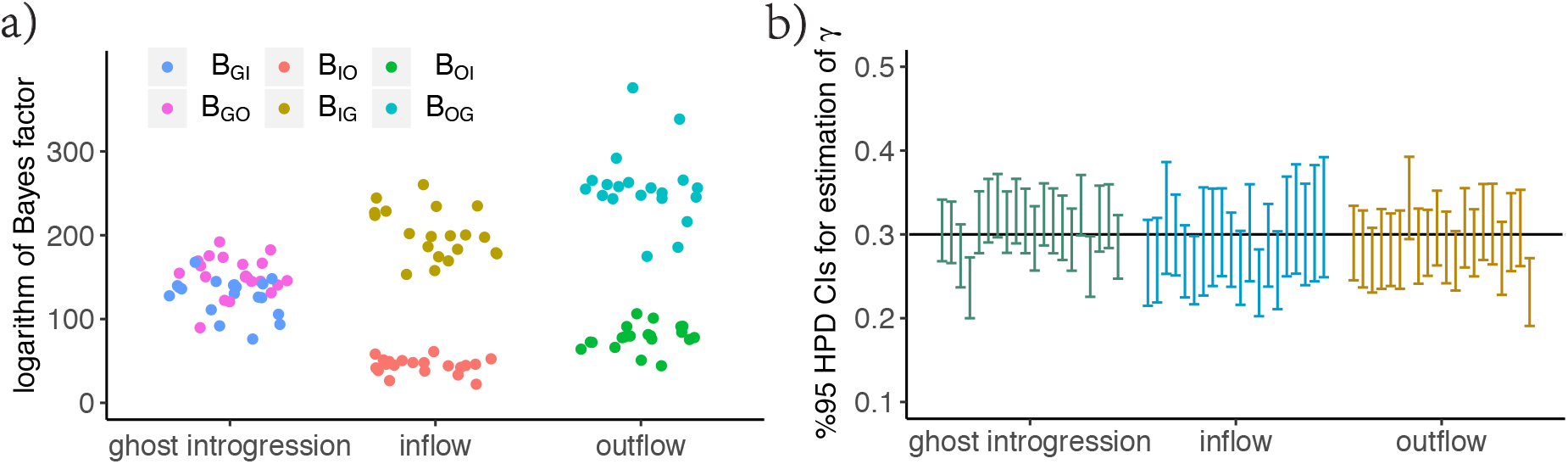
Results of the full-likelihood method BPP. a)The three introgression scenarios on x-axis are simulated based on the specific parameter combination (*C*_1_, *C*_2_, *γ*) = (0.3, 0.5, 0.3). For each simulated dataset, we computed Bayes factor to compare the three models of ghost introgression, inflow and outflow. The y-axis presents the logarithm of Bayes factors, representing the difference in logarithm of marginal likelihoods between the true model and two alternative false models. B_ij_ refers to the Bayes factor for model i against model j (G: ghost introgression, I: inflow, O: outflow). In all introgression scenarios, the Bayes factors strongly support the true model across 20 replicate datasets. b) The 95% highest-probability-density (HPD) credibility intervals (CIs) for the parameter *γ* under the true model, with the black line indicating the true value.

### Results of Ancient Introgression Scenarios

We also explored the performance of PhyloNet/MPL in detecting ancient ghost introgression. To do this, we simulated ancient introgression scenarios, wherein lineages undergo subsequent divergence events after an introgression event (Figs. 5a-c). We set the time intervals between the introgression event and the post-introgression divergence events (e.g., the divergence of *A*_*1*_ and *A*_*2*_) at 0.5 or 5 coalescent units to assess how this affects the performance of PhyloNet/MPL.

**Figure 5.**
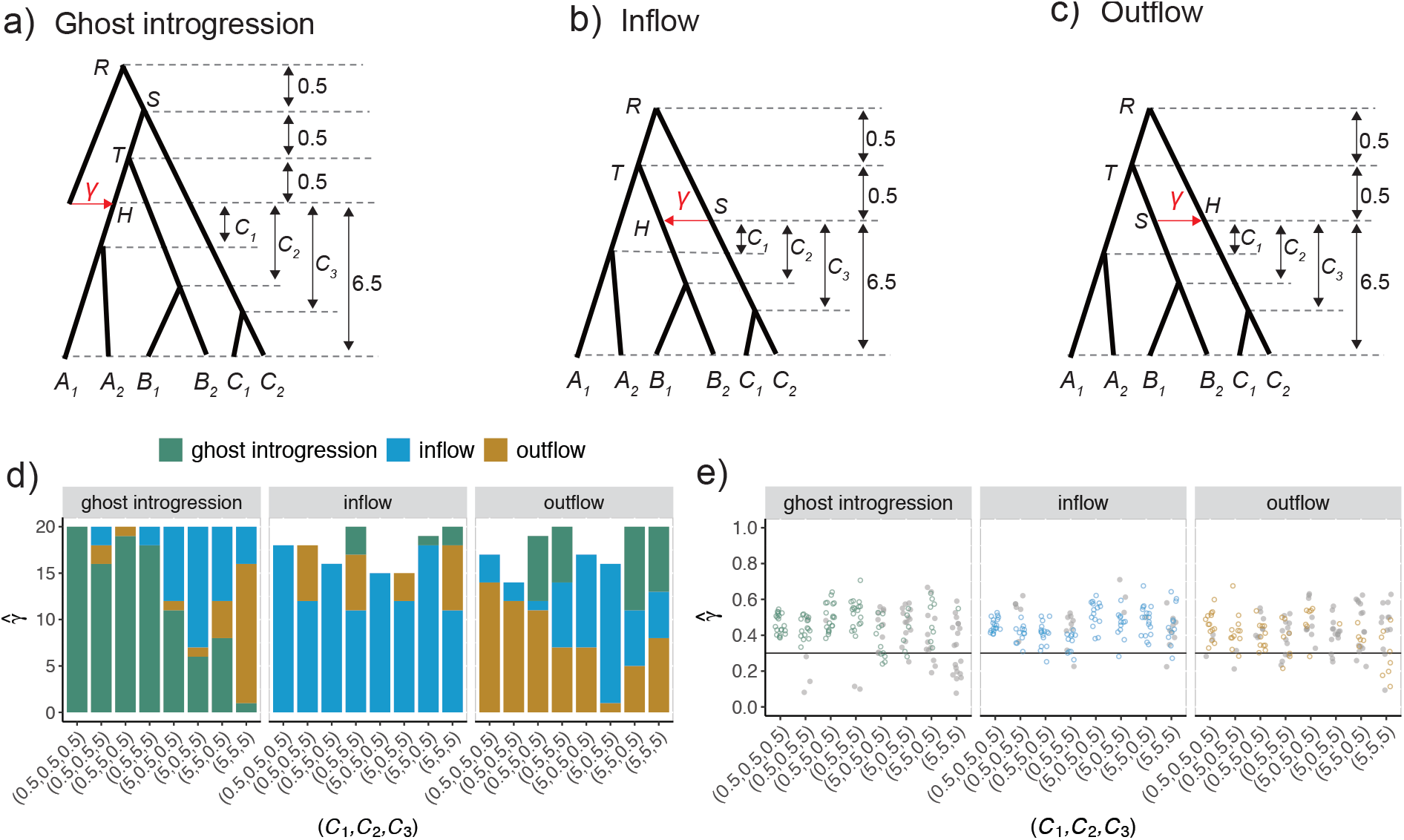
Results of PhyloNet/MPL in ancient introgression scenarios. a-c) The ancient scenarios of ghost introgression, inflow and outflow, along with the corresponding parameter settings. *C*_*i*_ (*i* = 1, 2, 3) denote the time intervals of the introgression event and the divergence event of species *A*_*1*_ and *A*_*2*_ or species *B*_*1*_ and *B*_*2*_ or species *C*_*1*_ and *C*_*2*_, respectively, and were set to 0.5 or 5 coalescent units. The introgression probability *γ* was fixed at 0.3. d-e) Plots of results for the network inference and *γ* estimation in simulations. The strips located on the top of each plot represent the true scenarios. The x-axis is labelled with the parameter combination (*C*_1_, *C*_2_, *C*_3_). d) Estimation of network topologies. The numbers of three inferred network topologies (i.e., topologies in a-c) among 20 replicates are displayed by corresponding colored bars. The remaining replicates that are not shown correspond to other inferred network topologies. e) Estimation of introgression probability *γ*. The solid line shows the true value. Colored and grey points represent the estimates in the true and other two false networks, respectively.

When the times between the introgression and divergence events are long, with (*C*_1_, *C*_2_, *C*_3_) = (5, 5, 5), the three networks that signify ghost introgression, inflow and outflow remain indistinguishable (Fig. 5d). When the introgression event is quickly followed by subsequent divergence events, with (*C*_1_, *C*_2_, *C*_3_) = (0.5, 0.5, 0.5), the networks can be inferred with 100% accuracy in the cases of ghost introgression and with at least 70% accuracy in inflow and outflow cases (Fig. 5d). Interestingly, *γ* tends to be overestimated in all scenarios and most parameter combinations (Fig. 5e).

## Analyses of Empirical Data: *J**altomata*

To investigate whether the above findings apply to real-world cases, we reanalyzed the transcriptome data from fourteen *Jaltomata* species (Wu et al. 2018b) and the outgroup *Solanum lycopersicum. Jaltomata* has experienced a recent radiation resulting in major subgroups with distinct mature fruit colours (Miller et al. 2011). Using the D-statistic, the original study by Wu et al. (2018b) reported an ancient introgression event between the early-diverging purple-fruited clade and the red-fruited lineage (*J. auriculata*). A recent study by Tiley et al. (2023) first estimated a species tree using ASTRAL (Zhang et al. 2018b) and then identified three introgression events in *Jaltomata* using SNaQ (Solís-Lemus and Ané 2016): an ancient ghost introgression event from an unknown ancestral lineage of *Jaltomata* to the common ancestor of the green- and orange-fruited clades, an inflow event between the extant lineages within the purple-fruited clade, and an outflow event from orange-fruited *J. umbellata* to the green-fruited common ancestor (Fig. 6a).

**Figure 6.**
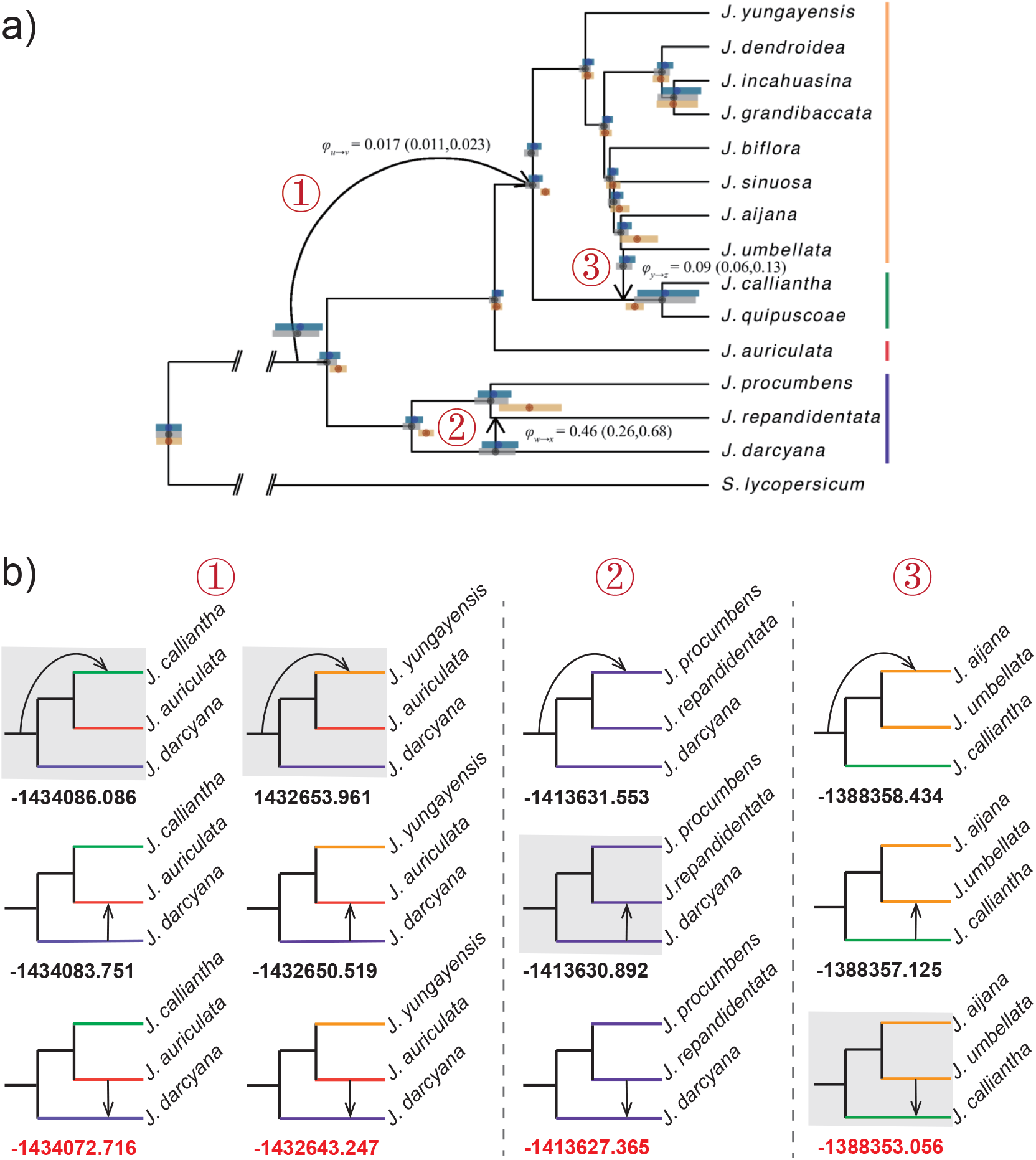
Inferred introgression events among *Jaltomata* species. a) The network topology inferred by SNaQ in Tiley et al. (2023; Figure 7) displays three introgression events. Vertical bars along the right show fruit colors among lineages. b) Comparison of ghost introgression, inflow and outflow models for each introgression event using BPP. The models labelled by the grey shading align with the network topology in Figure 6a. Each model is marked with its corresponding log margimal likelihood value below, with the red one indicating the best model.

Here, we revisit the three introgression events identified by the heuristic method SNaQ. We first investigated the ancient ghost introgression event from an unknown ancestral lineage of *Jaltomata* to the common ancestor of the green- and orange-fruited clades. We considered all possible combinations of species within the purple-fruited clade, the red-fruited clade, and the subgroup comprising the orange- and green-fruited clades, resulting in a total of 30 species trios (Supplementary Tables 1-2). HyDe and PhyloNet/MPL were used to reanalyze each species trio. For HyDe, the concatenated sequence alignments for every species trio, along with the outgroup *Solanum lycopersicum*, were used as input, and the *P*-value was calculated using the jackknife method. For PhyloNet/MPL, we estimated maximum likelihood (ML) gene trees from 6,431 one-to-one ortholog alignments using IQ-TREE v2 (Minh et al. 2020) with ModelFinder (Kalyaanamoorthy et al. 2017) to select the best-fitting substitution model and 1000 ultrafast bootstraps (-B 1000 -m MFP). These unrooted gene trees were then rooted with the outgroup *Solanum lycopersicum*. The subtrees for a focal species trio were extracted from the estimated gene trees as input for PhyloNet/MPL. Furthermore, due to the extensive computation required, BPP was used to analyze only two trios consisting of purple-fruited *J. darcyana*, red-fruited *J. auriculata*, and either green-fruited *J. calliantha* or orange-fruited *J. yungayensis*. To save computational time, we used only 1,000 orthologous alignments to compare the three models of ghost introgression, inflow, and outflow under the backbone topology from Tiley et al. (2023) by calculating the marginal likelihood values with 16 quadrature points. For each of the 16 MCMC runs, we used 50,000 iterations for the burnin, after which we took 1,000,000 posterior samples, sampling every 2 iterations. We then conducted the A00 analysis to estimate parameters for the optimal introgression model. Each run took ∼11 hrs using single threat. The inflow event and the outflow event in Figure 6a were investigated following the same procedure as just mentioned.

Our results on each of the three introgression events are presented in Figure 6 and Supplementary Tables 1-3. First, let us look at the ghost introgression event, labeled as ① in Fig. 6a. According to the HyDe analysis, the inflow scenario involving red-fruited *J. auriculata* as the hybrid was supported by all considered trios, with the estimated introgression probability varying from 0.0236 to 0.0526 across different species trios (Supplementary Table 1). However, PhyloNet/MPL failed to distinguish between the three scenarios of ghost introgression, inflow, and outflow as their log pseudo-likelihood values were close to each other (Supplementary Table 2). The introgression probability estimated by PhyloNet/MPL ranged widely from 0.075231 to 0.486673 across different trios. As expected, BPP was able to distinguish between the three introgression scenarios. For both trios (*J. darcyana, J. auriculata, J. calliantha*) and (*J. darcyana, J. auriculata, J. yungayensis*), the log marginal likelihood values for the outflow model were larger as compared to the ghost introgression and inflow models, indicating that the outflow model provided a better fit to the observed data (Fig. 6b). Our results are consistent with the ancient introgression between red-fruited *J. auriculata* and the purple-fruited clade identified by the D-statistic in the original study of Wu et al. (2018b) but contradict the conclusion of ghost introgression in Tiley et al. (2023). BPP estimated the introgression probability of the suppurted outflow event as 0.0584 and 0.05 for the two trios (Supplementary Table 3).

Next, we investigated the inflow event (labeled as ② in Fig. 6a) within the purple-fruited clade by considering the sole species trio of *J. procumbens, J. repandidentata*, and *J. darcyana*. The result of HyDe supported the inflow event with an introgression probability of 0.266 (Supplementary Table 1). Again, PhyloNet/MPL could not differentiate between the three scenarios of ghost introgression, inflow, and outflow with very close log pseudo-likelihood values of -6987.353797, -6987.352815, and -6987.353569, respectively (Supplementary Table 2). In contrast, BPP favored the outflow scenario with a log marginal likelihood of -1413627.365, while the ghost introgression and inflow models had lower log marginal likelihood values of -1413631.553 and -1413630.892, respectively (Fig. 6b). This BPP result also contradicts the previous conclusion of an inflow event in Tiley et al. (2023). The introgression probability for the supported outflow event is estimated as 0.11 (Supplementary Table 3).

Finally, we examined the outflow event (labeled as ③ in Fig. 6a) from orange-fruited *J. umbellata* to the green-fruited clade. We used HyDe and PhyloNet/MPL to analyze 14 different species trios, each involving *J. umbellata*, the green-fruited clade, and the orange-fruited clade. The HyDe results for all of these trios supported an inflow event with *J. umbellata* as the hybrid (Supplementary Table 1). PhyloNet/MPL still could not distinguish between the three scenarios (Supplementary Table 2). The trio including *J. calliantha, J. umbellata*, and *J. aijana* was analyzed in BPP. The outflow model was favored, with a logarithm of marginal likelihood of -1388353.056 compared to -1388358.434 for ghost introgression and -1388357.125 for inflow introgression (Fig. 6b). This confirms the original conclusion of Tiley et al. (2023). The introgression probability was estimated to be 0.081, close to the original estimate of 0.09 in Tiley et al. (2023) (Supplementary Table 3).

## Discussion

Ghost introgression has garnered significant attention among evolutionary biologists in recent years. The studies conducted by Pang and Zhang (2023) and Tiley et al. (2023) have established the significant impact of ghost introgression on species tree inference and divergence time estimation. Tricou et al. (2022b) demonstrated that ignoring ghost introgression can lead to the misidentification of both donors and recipients of introgression events using the D-statistic. In this article, we aim to determine the potential of currently popular phylogenetic tests of introgression in detecting ghost introgression and the pitfalls that may arise from failing to account for it.

### Unidentifiability of Ghost Introgression in Heuristic Methods

Based on theoretical analyses and simulation results, our study reveals that heuristic methods such as HyDe and PhyloNet/MPL cannot accurately identify ghost introgression even in the simplest cases of three focal species and a distant outgroup. This limitation arises from the fact that site patterns or gene tree topologies can only indicate whether introgression has occurred, but cannot differentiate between different types of introgression such as ghost introgression, inflow, or outflow. Although HyDe and PhyloNet/MPL face the same issue of network unidentifiability, they exhibit distinct patterns of behavior. Specifically, due to the assumption of hybrid speciation, HyDe’s interpretation is unequivocally oriented towards inflow cases when the statistically significant test confirms the presence of introgression. As such, HyDe works well only in inflow cases, but it can misidentify both the donor and recipient of introgression in both ghost introgression and outflow cases. By contrast, PhyloNet/MPL is liable to wrongly identify ghost introgression as ingroup introgressions and *vice versa*. Therefore, while HyDe and PhyloNet/MPL can provide indications of a specific introgression scenario, their conclusions should not be regarded as definitive. As with the D-statistic (Tricou et al. 2022b), the genuine interpretation of an introgression outcome when utilizing HyDe and PhyloNet/MPL should be a set of potential scenarios. That is, three scenarios including ghost introgression, inflow, and outflow, should be given equal consideration.

The introgression probability *γ* is unidentifiable when using HyDe and PhyloNet/MPL to detect ghost introgression. In HyDe, the accuracy of *γ* estimation is determined by the ratio of two internal branch lengths, i.e., *C*_*T*_ and *C*_*S*_ (or *l*_*T*_ and *l*_*S*_), which reflect the levels of ILS within the speciation and introgression histories, respectively (Fig. 2b). When *C*_*T*_ = *C*_*S*_, *γ* can be estimated with high accuracy, even when HyDe makes an erroneous inference about the introgression scenario. In contrast to HyDe, the estimates of *γ* by PhyloNet/MPL are more dispersed across multiple replicates. The degree of dispersion also depends on both branch lengths *C*_*T*_ and *C*_*S*_ (Fig. 2a). Longer branch lengths (i.e. weaker ILS) can lead to more precise estimates of *γ* in PhyloNet/MPL.

In ancient introgression scenarios where lineage divergence events occur after the introgression event (Figs. 5a-c), identifying ghost introgression presents similar challenges, especially when the time gap between introgression and subsequent divergence is significant. The challenges encountered by heuristic network inference methods can be attributed to the fact that sequences from recently diverged species have a high chance of coalescing within the long internal branch, making the introgression inference effectively the same as in the aforementioned three-taxon cases. Conversely, if the introgression event is followed immediately by the divergence events, it can result in a greater variety of gene tree topologies, providing the necessary information to accurately identify ghost introgression or introgressions between extant taxa (Zhu and Degnan 2017).

### The Promise of Full-likelihood Methods in Detecting Ghost Introgression

Several studies have highlighted the statistical advantages of using full-likelihood methods, as opposed to heuristic methods, in species tree inference, parameter estimation, and gene flow detection (Xu and Yang 2016; Flouri et al. 2020; Jiao et al. 2021; Zhu and Yang 2021; Ji et al. 2023). Ji et al. (2023) have shown that the full-likelihood method BPP is more efficient than the heuristic method HyDe for identifying gene flow, regardless of whether the involved species are sisters or not. On the other hand, our research focuses on exploring the advantages of full-likelihood methods in distinguishing different introgression scenarios that are compatible with a significant D-statistic.

Most satisfyingly, full-likelihood methods are capable of identifying ghost introgression. These methods take into account the information of topologies and branch lengths in gene trees, enabling discrimination between ghost introgression and introgressions between non-sister species that are indistinguishable by heuristic methods using topologies alone. The importance of branch lengths in network identifiability has been underscored in Yu et al. (2012) and Zhu and Degnan (2017) as well. Additionally, including branch lengths in gene trees can render some model parameters identifiable, such as population sizes, species divergence times, and introgression probabilities, which can help researchers better understand the evolutionary histories of species.

A major drawback of full-likelihood methods is their considerable computational burden, restricting their application to a limited number of taxa. Therefore, we recommend using full-likelihood methods only when putative introgression events have been implied by heuristic methods, just as what we have done in the above analysis of *Jaltomata*. Besides, the implementation of MSci in BPP assumes the molecular clock and is thus unsuitable for distantly related species that likely violate this assumption (Flouri et al. 2020). In contrast, heuristic methods make use of frequencies of gene tree topologies, thus are robust to the violation of the molecular clock assumption. The final caveat is that detecting ghost introgression necessitates a known species tree, just as most phylogenomic methods for introgression inference (Hibbins and Hahn 2022b). However, it can be challenging, if not impossible, to accurately infer the species tree from sequence data due to extensive introgression affecting much of the genome (Mallet et al. 2016; Jiao et al. 2020). An alternative approach may involve utilizing genome-structure information, such as gene order or content, which is more resistant to the effects of gene flow. Incorporating these features into the inference of species tree topologies could provide a promising avenue for overcoming this challenge (Zhao et al. 2021; Ding et al. 2023).

### Implications for Real Data Analysis of Ghost Introgression

The simulation studies conducted by Tricou et al. (2022b, 2022a) suggest that ghost introgressions are probably more likely to occur than ingroup introgressions. In recent years, empirical evidence for ghost introgression begins to appear in phylogenomic datasets such as those on *Jaltomata* species (Tiley et al. 2023) and on *Thuja* species (Li et al. 2022). But it is notable that most of such studies relied heavily on heuristic methods such as PhyloNet/MPL and SNaQ as they are computationally tractable for large-scale genomic data involving more than a few taxa. In the present study, we have unequivocally demonstrated that these heuristic methods usually confuse ghost introgression and introgression between non-sister species. This raises doubts about the biological conclusions regarding introgression events drawn from studies that rely solely on these heuristic methods.

Our reanalysis of the empirical dataset from *Jaltomata* species showcases the limitations of heuristic methods in distinguishing between ghost, inflow, and outflow introgressions and highlights the promise of full-likelihood methods for accurately inferring ghost introgression. In our reanalysis of the three introgression events among *Jaltomata* species inferred by SNaQ (Tiley et al. 2023), HyDe consistently identified them as inflow introgression, while PhyloNet/MPL could not distinguish between the three introgression scenarios with very close pseudo-likelihood values. These results align perfectly with our theoretical analyses and simulation results. By comparison, the full-likelihood BPP analysis supported an outflow event from the red-fruited *J. auriculata* to the purple-fruited clade, rather than the previously suggested ghost introgression in Tiley et al. (2023), as well as favored another outflow event from purple-fruited *J. repandidentata* to *J. arcyana* instead of the previously proposed inflow event in the reverse direction (Tiley et al. 2023). In other words, two of the three introgression events inferred by SNaQ are probably wrong.

We also reanalyzed the ghost introgression event among *Thuja* species as discovered by Li et al. (2022) (more details given in Supplementary Note 3 and Table S4). Our results, which were obtained using the selected species trio, indicated once again that HyDe and PhyloNet/MPL performed poorly, while BPP provided strong evidence in favor of the ghost introgression model, a conclusion also drawn from analyzing all *Thuja* species using PhyloNet/MPL in the original study. This firmly establishes the presence of ghost introgression among *Thuja* species, and we expect more properly confirmed cases of ghost introgression to come out in future.

Here, we propose using the full-likelihood method BPP as a crucial step in validating gene flow events identified by heuristic methods. This involves comparing the focal gene flow event with two alternative introgression models through marginal likelihood calculation for relevant species trios, as demonstrated in our analyses of *Jaltomata* and *Thuja*. We have demonstrated that this strategy is both computationally feasible and effective in identifying ghost introgression. We recommend exercising caution when inferring introgression events using heuristic methods alone and suggest adopting a full-likelihood approach whenever possible to confidently detect ghost introgression in empirical studies.

## Conclusion

Our study highlights the issue of non-identifiability of ghost introgression in heuristic methods such as HyDe, PhyloNet/MPL or SNaQ, which raises doubts about the reliability of previous introgression conclusions that heavily or solely rely on these methods for inferring gene flow events. Therefore, we caution against interpreting heuristic method results without considering the possibility of ghost introgression and recommend adopting full-likelihood methods to make further judgments about the nature of the introgression event. This strategy is both computationally feasible and effective, making it the best practice for detecting ghost introgression in empirical studies. Undoubtedly, future efforts should be directed towards improving the statistical efficiency of heuristic methods and the computational efficiency of Bayesian MCMC algorithms employed in full-likelihood methods.

## Acknowledgements

We are grateful to Susanne Renner for her encouragement and constructive comments. We would like to thank Ya-Mei Ding, Wei-Ning Bai, Erli Pang, and Yan-Ping Guo for helpful discussions.

## Funding

This work was supported by the National Natural Science Foundation of China (31421063 and 32170223), the “111” Program of Introducing Talents of Discipline to Universities (B13008), Beijing Advanced Innovation Program for Land Surface Processes, and the National Key R&D Program of China (2017YFA0605104).

## Notes

### Competing Interest Statement

The authors have declared no competing interest.

## References

Ai H, Fang X, Yang B, Huang Z, Chen H, Mao L, Zhang F, Zhang L, Cui L, He W, Yang J, Yao X, Zhou L, Han L, Li J, Sun S, Xie X, Lai B, Su Y, Lu Y, Yang H, Huang T, Deng W, Nielsen R, Ren J, Huang L. 2015. Adaptation and possible ancient interspecies introgression in pigs identified by whole-genome sequencing. Nat. Genet. 47:217–225.

Allman ES, Baños H, Rhodes JA. 2019. NANUQ: A method for inferring species networks from gene trees under the coalescent model. Algorithms for Molecular Biology 14:1–25.

Blischak PD, Chifman J, Wolfe AD, Kubatko LS. 2018. HyDe: A python package for genomescale hybridization detection. Syst. Biol. 67:821–829.

Ding Y-M, Cao Y, Zhang W-P, Chen J, Liu J, Li P, Renner SS, Zhang D-Y, Bai W-N. 2022. Population-genomic analyses reveal bottlenecks and asymmetric introgression from Persian into iron walnut during domestication. Genome Biol. 23:1–18.

Ding Y-M, Pang X-X, Cao Y, Zhang W-P, Renner SS, Zhang D-Y, Bai W-N. 2023. Genome structure-based Juglandaceae phylogenies contradict alignment-based phylogenies and substitution rates vary with DNA repair genes. Nat. Commun. 14:617.

Edelman NB, Mallet J. 2021. Prevalence and adaptive impact of introgression. Annu. Rev. Genet. 55:265–283.

Esquerré D, Keogh JS, Demangel D, Morando M, Avila LJ, Sites JW, Jr., Ferri-Yáñez F, Leaché AD. 2021. Rapid radiation and rampant reticulation: Phylogenomics of South American Liolaemus lizards. Syst. Biol. 71:286–300.

Figueiró HV, Li G, Trindade FJ, Assis J, Pais F, Fernandes G, Santos SH, Hughes GM, Komissarov A, Antunes A. 2017. Genome-wide signatures of complex introgression and adaptive evolution in the big cats. Sci. Adv. 3:e1700299.

Flouri T, Jiao XY, Rannala B, Yang ZH. 2020. A Bayesian implementation of the multispecies coalescent model with introgression for phylogenomic analysis. Mol. Biol. Evol. 37:1211–1223.

Fontaine MC, Pease JB, Steele A, Waterhouse RM, Neafsey DE, Sharakhov IV, Jiang X, Hall AB, Catteruccia F, Kakani E, Mitchell SN, Wu Y-C, Smith HA, Love RR, Lawniczak MK, Slotman MA, Emrich SJ, Hahn MW, Besansky NJ. 2015. Extensive introgression in a malaria vector species complex revealed by phylogenomics. Science 347:1258524–1258524.

Green RE, Krause J, Briggs AW, Maricic T, Stenzel U, Kircher M, Patterson N, Li H, Zhai W, Fritz MH-Y. 2010. A draft sequence of the Neandertal genome. Science 328:710–722.

Hey J, Chung Y, Sethuraman A, Lachance J, Tishkoff S, Sousa VC, Wang Y. 2018. Phylogeny estimation by integration over isolation with migration models. Mol. Biol. Evol. 35:2805– 2818.

Hibbins MS, Hahn MW. 2022a. Distinguishing between histories of speciation and introgression using genomic data. bioRxiv doi: https://doi.org/10.1101/2022.09.07.506990.

Hibbins MS, Hahn MW. 2022b. Phylogenomic approaches to detecting and characterizing introgression. Genetics 220.

Huang J, Flouri T, Yang Z. 2020. A simulation study to examine the information content in phylogenomic data sets under the multispecies coalescent model. Mol. Biol. Evol. 37:3211–3224.

Hudson RR. 2002. Generating samples under a Wright-Fisher neutral model of genetic variation. Bioinformatics 18:337–338.

Ji J, Jackson DJ, Leaché AD, Yang Z. 2023. Power of Bayesian and heuristic tests to detect cross-species introgression with reference to gene flow in the Tamias quadrivittatus group of North American chipmunks. Syst. Biol. doi: https://doi.org/10.1093/sysbio/syac077.

Jiao X, Flouri T, Rannala B, Yang Z. 2020. The impact of cross-species gene flow on species tree estimation. Syst. Biol. 69:830–847.

Jiao X, Flouri T, Yang Z. 2021. Multispecies coalescent and its applications to infer species phylogenies and cross-species gene flow. Natl. Sci. Rev. 8:wab127.

Jones MR, Mills LS, Alves PC, Callahan CM, Alves JM, Lafferty DJ, Jiggins FM, Jensen JD, Melo-Ferreira J, Good JM. 2018. Adaptive introgression underlies polymorphic seasonal camouflage in snowshoe hares. Science 360:1355–1358.

Kalyaanamoorthy S, Minh BQ, Wong TK, Von Haeseler A, Jermiin LS. 2017. Modelfinder: Fast model selection for accurate phylogenetic estimates. Nat. Methods 14:587–589.

Kong S, Kubatko LS. 2021. Comparative performance of popular methods for hybrid detection using genomic data. Syst. Biol. 70:891–907.

Kubatko LS, Chifman J. 2019. An invariants-based method for efficient identification of hybrid species from large-scale genomic data. BMC Evol. Biol. 19:1–13.

Kuhlwilm M, Han S, Sousa VC, Excoffier L, Marques-Bonet T. 2019. Ancient admixture from an extinct ape lineage into bonobos. Nat. Ecol. Evol. 3:957–965.

Lartillot N, Philippe H. 2006. Computing Bayes factors using thermodynamic integration. Syst. Biol. 55:195–207.

Li J, Zhang Y, Ruhsam M, Milne RI, Wang Y, Wu D, Jia S, Tao T, Mao K. 2022. Seeing through the hedge: Phylogenomics of Thuja (Cupressaceae) reveals prominent incomplete lineage sorting and ancient introgression for Tertiary relict flora. Cladistics 38:187–203.

Mallet J, Besansky N, Hahn MW. 2016. How reticulated are species? Bioessays 38:140–149.

Meleshko O, Martin MD, Korneliussen TS, Schröck C, Lamkowski P, Schmutz J, Healey A, Piatkowski BT, Shaw AJ, Weston DJ, Flatberg KI, Szövényi P, Hassel K, Stenøien HK. 2021. Extensive genome-wide phylogenetic discordance is due to incomplete lineage sorting and not ongoing introgression in a rapidly radiated bryophyte genus. Mol. Biol. Evol. 38:2750–2766.

Mendes FK, Hahn MW. 2018. Why concatenation fails near the anomaly zone. Syst. Biol. 67:158–169.

Miller RJ, Mione T, Phan H-L, Olmstead RG. 2011. Color by numbers: Nuclear gene phylogeny of Jaltomata (Solanaceae), sister genus to Solanum, supports three clades differing in fruit color. Syst. Bot. 36:153–162.

Minh BQ, Schmidt HA, Chernomor O, Schrempf D, Woodhams MD, von Haeseler A, Lanfear R. 2020. IQ-TREE 2: New models and efficient methods for phylogenetic inference in the genomic era. Mol. Biol. Evol. 37:1530–1534.

Ottenburghs J. 2020. Ghost introgression: Spooky gene flow in the distant past. Bioessays 42:e2000012.

Pang X-X, Zhang D-Y. 2023. Impact of ghost introgression on coalescent-based species tree inference and estimation of divergence time. Syst. Biol. doi: https://doi.org/10.1093/sysbio/syac047.

Pardi F, Scornavacca C. 2015. Reconstructible phylogenetic networks: Do not distinguish the indistinguishable. PLoS Comp. Biol. 11:e1004135.

Rambaut A, Grass NC. 1997. Seq-Gen: An application for the Monte Carlo simulation of DNA sequence evolution along phylogenetic trees. Bioinformatics 13:235–238.

Rannala B, Yang Z. 2017. Efficient Bayesian species tree inference under the multispecies coalescent. Syst. Biol. 66:823–842.

Rocha JL, Vaz Pinto P, Siegismund HR, Meyer M, Jansen van Vuuren B, Veríssimo L, Ferrand N, Godinho R. 2022. African climate and geomorphology drive evolution and ghost introgression in sable antelope. Mol. Ecol. 31:2968–2984.

Sankararaman S, Mallick S, Dannemann M, Prüfer K, Kelso J, Pääbo S, Patterson N, Reich D. 2014. The genomic landscape of Neanderthal ancestry in present-day humans. Nature 507:354–357.

Solís-Lemus C, Ané C. 2016. Inferring phylogenetic networks with maximum pseudolikelihood under incomplete lineage sorting. PLoS Genet. 12:e1005896.

Suvorov A, Kim BY, Wang J, Armstrong EE, Peede D, D’agostino ER, Price DK, Waddell PJ, Lang M, Courtier-Orgogozo V. 2022. Widespread introgression across a phylogeny of 155 Drosophila genomes. Curr. Biol. 32:111–123.

Taylor SA, Larson EL. 2019. Insights from genomes into the evolutionary importance and prevalence of hybridization in nature. Nat. Ecol. Evol. 3:170–177.

Tiley GP, Flouri T, Jiao X, Poelstra JW, Xu B, Zhu T, Rannala B, Yoder AD, Yang Z. 2023. Estimation of species divergence times in presence of cross-species gene flow. Syst. Biol. doi: https://doi.org/10.1093/sysbio/syad015.

Tricou T, Tannier E, de Vienne DM. 2022a. Ghost lineages can invalidate or even reverse findings regarding gene flow. PLoS Biol. 20:e3001776.

Tricou T, Tannier E, de Vienne DM. 2022b. Ghost lineages highly influence the interpretation of introgression tests. Syst. Biol. 71:1147–1158.

Wang M, Zhang L, Zhang Z, Li M, Wang D, Zhang X, Xi Z, Keefover-Ring K, Smart LB, DiFazio SP, Olson MS, Yin T, Liu J, Ma T. 2020. Phylogenomics of the genus Populus reveals extensive interspecific gene flow and balancing selection. New Phytol. 225:1370– 1382.

Wen D, Nakhleh L. 2018. Coestimating reticulate phylogenies and gene trees from multilocus sequence data. Syst. Biol. 67:439–457.

Wu D-D, Ding X-D, Wang S, Wójcik JM, Zhang Y, Tokarska M, Li Y, Wang M-S, Faruque O, Nielsen R, Zhang Q, Zhang Y-P. 2018a. Pervasive introgression facilitated domestication and adaptation in the Bos species complex. Nat. Ecol. Evol. 2:1139–1145.

Wu M, Kostyun JL, Hahn MW, Moyle LC. 2018b. Dissecting the basis of novel trait evolution in a radiation with widespread phylogenetic discordance. Mol. Ecol. 27:3301–3316.

Xu B, Yang Z. 2016. Challenges in species tree estimation under the multispecies coalescent model. Genetics 204:1353–1368.

Yang W, Feiner N, Pinho C, While GM, Kaliontzopoulou A, Harris DJ, Salvi D, Uller T. 2021. Extensive introgression and mosaic genomes of Mediterranean endemic lizards. Nat. Commun. 12:2762.

Yang Z. 2015. The BPP program for species tree estimation and species delimitation. Curr Zool 61:854–865.

Yang Z, Flouri T. 2022. Estimation of cross-species introgression rates using genomic data despite model unidentifiability. Mol. Biol. Evol. 39:msac083.

Yu Y, Degnan JH, Nakhleh L. 2012. The probability of a gene tree topology within a phylogenetic network with applications to hybridization detection. PLoS Genet. 8:456– 465.

Yu Y, Dong J, Liu KJ, Nakhleh L. 2014. Maximum likelihood inference of reticulate evolutionary histories. Proc. Natl. Acad. Sci. USA 111:16448–16453.

Yu Y, Nakhleh L. 2015. A maximum pseudo-likelihood approach for phylogenetic networks. BMC Genom. 16:S10.

Zhang BW, Xu LL, Li N, Yan PC, Jiang XH, Woeste KE, Lin K, Renner SS, Zhang DY, Bai WN. 2019. Phylogenomics reveals an ancient hybrid origin of the Persian walnut. Mol. Biol. Evol. 36:2451–2461.

Zhang C, Ogilvie HA, Drummond AJ, Stadler T. 2018a. Bayesian inference of species networks from multilocus sequence data. Mol. Biol. Evol. 35:504–517.

Zhang C, Rabiee M, Sayyari E, Mirarab S. 2018b. ASTRAL-III: Polynomial time species tree reconstruction from partially resolved gene trees. BMC Bioinform. 19:153.

Zhao T, Zwaenepoel A, Xue J-Y, Kao S-M, Li Z, Schranz ME, Van de Peer Y. 2021. Wholegenome microsynteny-based phylogeny of angiosperms. Nat. Commun. 12:3498.

Zhu S, Degnan JH. 2017. Displayed trees do not determine distinguishability under the network multispecies coalescent. Syst. Biol. 66:283–298.

Zhu T, Yang Z. 2021. Complexity of the simplest species tree problem. Mol. Biol. Evol. 38:3993–4009.

