## supplemental materials for "Detection of Ghost Introgression from Phylogenomic Data Requires a Full-Likelihood Approach"

**Supplementary materials to Pang and Zhang's manuscript entitled "Detection of Ghost Introgression from Phylogenomic Data Requires a Full-Likelihood Approach"**

### Supplementary Note 1

To calculate the frequency of each site pattern per nucleotide site in the three introgression scenarios in Figures 1a-c, we weight the internal branch lengths of the relevant gene trees by their probabilities and sum them across coalescent histories.

Ghost introgression scenario is shown in Figure 1a. Following the speciation history of $AB|C$ with a probability of $1-\gamma$, the gene tree $G_1$ arises if sequences *a* and *b* coalesce in species *T*. The coalescent time *t* ($\tau_T < t < \tau_S$) of sequences *a* and *b* coalescing in species *T* has the probability density of $(1-\gamma)\frac{2}{\theta_T}e^{-\frac{2}{\theta_T}(t-\tau_T)}$, corresponding to the internal branch length in $G_1$ with $\tau_S - t + (1-e^{-\frac{2l_S}{\theta_S}})\frac{\theta_S}{2} + e^{-\frac{2l_S}{\theta_S}}\frac{\theta_R}{2}$. Thus, the sum of the internal branches of this part of gene trees $G_1$ is $\int_{\tau_T}^{\tau_S}(1-\gamma)\frac{2}{\theta_T}e^{-\frac{2(t-\tau_T)}{\theta_T}} \times (\tau_S - t + (1-e^{-\frac{2l_S}{\theta_S}})\frac{\theta_S}{2} + e^{-\frac{2l_S}{\theta_S}}\frac{\theta_R}{2})\,dt = (1-\gamma)(l_T + (1-e^{-\frac{2l_T}{\theta_T}})\frac{(1-e^{-\frac{2l_S}{\theta_S}})\theta_S + e^{-\frac{2l_S}{\theta_S}}\theta_R - \theta_T}{2})$. Otherwise, both coalescence events of all three sequences occur in the species *S* or *R*, producing the sum of internal branch lengths of $\frac{1}{3}(1-\gamma)e^{-\frac{2l_T}{\theta_T}}\{\frac{\theta_S}{2}(1-e^{-\frac{6l_S}{\theta_S}} - \frac{3}{8}(1-e^{-\frac{4l_S}{\theta_S}})) + \frac{\theta_R}{2}(e^{-\frac{6l_S}{\theta_S}} + \frac{3}{8}(1-e^{-\frac{4l_S}{\theta_S}}))\}$ for each gene tree. Following the introgressed history of $A|BC$ with a probability of $\gamma$, gene tree $G_2$ arises when sequences *b* and *c* coalesce in species *S*. The coalescent time *t* ($\tau_S < t < \tau_R$) of sequences *b* and *c* coalescing in species *S* has the probability density of $\gamma\frac{2}{\theta_S}e^{-\frac{2(t-\tau_S)}{\theta_S}}$, corresponding to the internal branch length of $\tau_R - t + \frac{\theta_R}{2}$. The sum of internal branches of the part of gene trees $G_2$ is $\int_{\tau_S}^{\tau_R}\gamma\frac{2}{\theta_S}e^{-\frac{2(t-\tau_S)}{\theta_S}} \times (\tau_R - t + \frac{\theta_R}{2})dt = \gamma(l_S + (1-e^{-\frac{2l_S}{\theta_S}})\frac{\theta_R-\theta_S}{2})$. Otherwise, all three sequences coalesce after entering species *R*, producing the sum of internal branch lengths of $\frac{\theta_R}{6}\gamma e^{-\frac{2l_S}{\theta_S}}$ for each gene tree. Thus, let $P = \frac{\theta_S}{2}(1 -$

$e^{-\frac{6l_S}{\theta_S}}-\frac{3}{8}(1-e^{-\frac{4l_S}{\theta_S}}))+\frac{\theta_R}{2}(e^{-\frac{6l_S}{\theta_S}}+\frac{3}{8}(1-e^{-\frac{4l_S}{\theta_S}}))$, there is the expression for the frequency of each site pattern per nucleotide site:

$$
\begin{aligned}
f(\text{ABAB}) &= \frac{\theta_R}{6}\gamma e^{-\frac{2l_S}{\theta_S}}+\frac{1}{3}(1-\gamma)e^{-\frac{2l_T}{\theta_T}}P \\
f(\text{ABBA}) &= (1-\gamma)(l_T+(1-e^{-\frac{2l_T}{\theta_T}})\frac{(1-e^{-\frac{2l_S}{\theta_S}})\theta_S+e^{-\frac{2l_S}{\theta_S}}\theta_R-\theta_T}{2})+ \\
&\quad \frac{\theta_R}{6}\gamma e^{-\frac{2l_S}{\theta_S}}+\frac{1}{3}(1-\gamma)e^{-\frac{2l_T}{\theta_T}}P \\
&= (1-\gamma)(l_T+(1-e^{-\frac{2l_T}{\theta_T}})\frac{(1-e^{-\frac{2l_S}{\theta_S}})\theta_S+e^{-\frac{2l_S}{\theta_S}}\theta_R-\theta_T}{2})+P(\text{ABAB}) \qquad (1) \\
f(\text{AABB}) &= \gamma(l_S+(1-e^{-\frac{2l_S}{\theta_S}})\frac{\theta_R-\theta_S}{2})+\frac{\theta_R}{6}\gamma e^{-\frac{2l_S}{\theta_S}}+\frac{1}{3}(1-\gamma)e^{-\frac{2l_T}{\theta_T}}P \\
&= \gamma(l_S+(1-e^{-\frac{2l_S}{\theta_S}})\frac{\theta_R-\theta_S}{2})+P(\text{ABAB})
\end{aligned}
$$

For the inflow case in Figure 1b, following the speciation history of $AB|C$ with a probability of $1-\gamma$, the gene tree $G_1$ arises if sequences *a* and *b* coalesce in species *T*. The coalescent time *t* ($\tau_T < t < \tau_R$) of sequences *a* and *b* coalescing in species *T* has the probability density of $(1-\gamma)\frac{2}{\theta_T}e^{-\frac{2}{\theta_T}(t-\tau_T)}$, corresponding to the internal branch length in $G_1$ with $\tau_S - t + \frac{\theta_R}{2}$. Thus, the sum of the internal branches of this part of gene trees $G_1$ is $\int_{\tau_T}^{\tau_R}(1-\gamma)\frac{2}{\theta_T}e^{-\frac{2(t-\tau_T)}{\theta_T}}\times(\tau_S-t+\frac{\theta_R}{2})\,dt=(1-\gamma)(l_T+(1-e^{-\frac{2l_T}{\theta_T}})\frac{\theta_R-\theta_T}{2})$. Otherwise, both coalescence events of all three sequences occur in the species *R*, producing the sum of internal branch lengths of $\frac{\theta_R}{6}(1-\gamma)e^{-\frac{2l_T}{\theta_T}}$ for each gene tree. Following the introgressed history of $A|BC$ with a probability of $\gamma$, gene tree $G_2$ arises when sequences *b* and *c* coalesce in species *S*. The coalescent time *t* ($\tau_S < t < \tau_R$) of sequences *b* and *c* coalescing in species *S* has the probability density of $\gamma\frac{2}{\theta_S}e^{-\frac{2(t-\tau_S)}{\theta_S}}$, corresponding to the internal branch length of $\tau_R - t + \frac{\theta_R}{2}$. The sum of internal branches of the part of gene trees $G_2$ is

$\int_{\tau_S}^{\tau_R}\gamma\frac{2}{\theta_S}e^{-\frac{2(t-\tau_S)}{\theta_S}}\times(\tau_R-t+\frac{\theta_R}{2})dt=\gamma(l_S+(1-e^{-\frac{2l_S}{\theta_S}})\frac{\theta_R-\theta_S}{2})$. Otherwise, all three sequences coalesce after entering species *R*, producing the sum of internal branch lengths of $\frac{\theta_R}{6}\gamma e^{-\frac{2l_S}{\theta_S}}$ for each gene tree. Thus, the expression for the frequency of each site pattern per nucleotide site is:

$$\begin{aligned}
f(\text{ABAB}) &= \frac{\theta_R}{6}(1-\gamma)e^{-\frac{2l_T}{\theta_T}}+\frac{\theta_R}{6}\gamma e^{-\frac{2l_S}{\theta_S}} \\
f(\text{ABBA}) &= (1-\gamma)(l_T+(1-e^{-\frac{2l_T}{\theta_T}})\frac{\theta_R-\theta_T}{2})+\frac{\theta_R}{6}(1-\gamma)e^{-\frac{2l_T}{\theta_T}}+\frac{\theta_R}{6}\gamma e^{-\frac{2l_S}{\theta_S}} \\
&= (1-\gamma)(l_T+(1-e^{-\frac{2l_T}{\theta_T}})\frac{\theta_R-\theta_T}{2})+P(\text{ABAB}) \\
f(\text{AABB}) &= \gamma(l_S+(1-e^{-\frac{2l_S}{\theta_S}})\frac{\theta_R-\theta_S}{2})+\frac{\theta_R}{6}(1-\gamma)e^{-\frac{2l_T}{\theta_T}}+\frac{\theta_R}{6}\gamma e^{-\frac{2l_S}{\theta_S}} \\
&= \gamma(l_S+(1-e^{-\frac{2l_S}{\theta_S}})\frac{\theta_R-\theta_S}{2})+P(\text{ABAB})
\end{aligned} \tag{2}$$

For the outflow introgression scenarios in Figure 1c, following the speciation history of $AB|C$ with a probability of $1-\gamma$, the gene tree $G_1$ arises if sequences *a* and *b* coalesce in species *T*. The coalescent time *t* ($\tau_T < t < \tau_R$) of sequences *a* and *b* coalescing in species *T* has the probability density of $(1-\gamma)\frac{2}{\theta_T}e^{-\frac{2}{\theta_T}(t-\tau_T)}$, corresponding to the internal branch length in $G_1$ with $\tau_S - t + \frac{\theta_R}{2}$. Thus, the sum of the internal branches of this part of gene trees $G_1$ is $\int_{\tau_T}^{\tau_S}(1-\gamma)\frac{2}{\theta_T}e^{-\frac{2(t-\tau_T)}{\theta_T}}\times(\tau_S-t+\frac{\theta_R}{2})\,dt=(1-\gamma)(l_T+(1-e^{-\frac{2l_T}{\theta_T}})\frac{\theta_R-\theta_T}{2})$. Otherwise, both coalescence events of all three sequences occur in the species *R*, producing the sum of internal branch lengths of $\frac{\theta_R}{6}(1-\gamma)e^{-\frac{2l_T}{\theta_T}}$ for each gene tree. Following the introgression history of $A|BC$ with a probability of $\gamma$, gene tree $G_2$ arises when sequences *b* and *c* coalesce in species *S*. The coalescent time *t* ($\tau_S < t < \tau_\mathrm{T}$) of sequences *b* and *c* coalescing in species *S* has the probability density of $\gamma\frac{2}{\theta_S}e^{-\frac{2(t-\tau_S)}{\theta_S}}$, corresponding to the internal branch length of $\tau_\mathrm{T}-t+(1-e^{-\frac{2l_T}{\theta_T}})\theta_T+e^{-\frac{2l_T}{\theta_T}}\theta_R$. The sum of internal branches of the part of gene trees $G_2$ is $\int_{\tau_S}^{\tau_\mathrm{T}}\gamma\frac{2}{\theta_S}e^{-\frac{2(t-\tau_S)}{\theta_S}}\times(\tau_\mathrm{T}-t+(1-e^{-\frac{2l_T}{\theta_T}})\theta_T+e^{-\frac{2l_T}{\theta_T}}\theta_R)dt=\gamma(l_S+(1-e^{-\frac{2l_S}{\theta_S}})\frac{(1-e^{-\frac{2l_T}{\theta_T}})\theta_T+e^{-\frac{2l_T}{\theta_T}}\theta_R-\theta_S}{2})$. Otherwise, all three sequences coalesce after entering species *T* or *R*, producing the sum of internal branch lengths of $\frac{1}{3}\gamma e^{-\frac{2l_S}{\theta_S}}\{\frac{\theta_T}{2}(1-e^{-\frac{6l_T}{\theta_T}}-\frac{3}{8}(1-e^{-\frac{4l_T}{\theta_T}}))+\frac{\theta_R}{2}(e^{-\frac{6l_T}{\theta_T}}+\frac{3}{8}(1-e^{-\frac{4l_T}{\theta_T}}))\}$ for each gene tree. Thus, let $P=\frac{\theta_T}{2}(1-e^{-\frac{6l_T}{\theta_T}}-\frac{3}{8}(1-e^{-\frac{4l_T}{\theta_T}}))+\frac{\theta_R}{2}(e^{-\frac{6l_T}{\theta_T}}+\frac{3}{8}(1-e^{-\frac{4l_T}{\theta_T}}))$, there is the expression for the frequency of each site pattern per nucleotide site:

$$
\begin{aligned}
f(ABAB) &= \frac{\theta_R}{6}(1-\gamma)e^{-\frac{2l_T}{\theta_T}}+\frac{1}{3}\gamma e^{-\frac{2l_S}{\theta_S}}P \\
f(ABBA) &= (1-\gamma)(l_T+(1-e^{-\frac{2l_T}{\theta_T}})\frac{\theta_R-\theta_T}{2}+\frac{\theta_R}{6}(1-\gamma)e^{-\frac{2l_T}{\theta_T}}+\frac{1}{3}\gamma e^{-\frac{2l_S}{\theta_S}}P \\
&= (1-\gamma)(l_T+(1-e^{-\frac{2l_T}{\theta_T}})\frac{\theta_R-\theta_T}{2})+P(ABAB) \\
f(AABB) &= \gamma(l_S+(1-e^{-\frac{2l_S}{\theta_S}})\frac{(1-e^{-\frac{2l_T}{\theta_T}})\theta_T+e^{-\frac{2l_T}{\theta_T}}\theta_R-\theta_S}{2})+\frac{\theta_R}{6}(1-\gamma)e^{-\frac{2l_T}{\theta_T}}+\frac{1}{3}\gamma e^{-\frac{2l_S}{\theta_S}}P \\
&= \gamma(l_S+(1-e^{-\frac{2l_S}{\theta_S}})\frac{(1-e^{-\frac{2l_T}{\theta_T}})\theta_T+e^{-\frac{2l_T}{\theta_T}}\theta_R-\theta_S}{2})+P(ABAB)
\end{aligned}
\qquad (3)
$$

**Supplementary Note 2**

Let $P_T = e^{-\frac{2l_T}{\theta_T}}$ and $P_S = e^{-\frac{2l_S}{\theta_S}}$. In the case of ghost introgression shown in Figure 1a, the density distribution of coalescent times of each sequence pair can be expressed as:

$$f(t_{ab}) = \begin{cases} (1-\gamma)\dfrac{2}{\theta_T}e^{-\frac{2}{\theta_T}(t_{ab}-\tau_T)}, & \tau_T < t_{ab} < \tau_S \\ (1-\gamma)P_T\dfrac{2}{\theta_S}e^{-\frac{2}{\theta_S}(t_{ab}-\tau_S)}, & \tau_S < t_{ab} < \tau_R \\ [(1-\gamma)P_T P_S + \gamma]\dfrac{2}{\theta_R}e^{-\frac{2}{\theta_R}(t_{ab}-\tau_R)}, & t_{ab} > \tau_R \end{cases}$$

$$f(t_{bc}) = \begin{cases} \dfrac{2}{\theta_S}e^{-\frac{2}{\theta_S}(t_{bc}-\tau_S)}, & \tau_S < t_{bc} < \tau_R \\ P_S\dfrac{2}{\theta_R}e^{-\frac{2}{\theta_R}(t_{bc}-\tau_R)}, & t_{bc} > \tau_R \end{cases} \tag{4}$$

$$f(t_{ac}) = \begin{cases} (1-\gamma)\dfrac{2}{\theta_S}e^{-\frac{2}{\theta_S}(t_{ac}-\tau_S)}, & \tau_S < t_{ac} < \tau_R \\ [(1-\gamma)P_S + \gamma]\dfrac{2}{\theta_R}e^{-\frac{2}{\theta_R}(t_{ac}-\tau_R)}, & t_{ac} > \tau_R \end{cases}$$

In the case of inflow shown in Figure 1b, the density distribution of coalescent times of each sequence pair is:

$$f(t_{ab}) = \begin{cases} (1-\gamma)\dfrac{2}{\theta_T}e^{-\frac{2}{\theta_T}(t_{ab}-\tau_T)}, & \tau_T < t_{ab} < \tau_R \\ [(1-\gamma)P_T + \gamma]\dfrac{2}{\theta_R}e^{-\frac{2}{\theta_R}(t_{ab}-\tau_R)}, & t_{ab} > \tau_R \end{cases}$$

$$f(t_{bc}) = \begin{cases} \gamma\dfrac{2}{\theta_S}e^{-\frac{2}{\theta_S}(t_{bc}-\tau_S)}, & \tau_S < t_{bc} < \tau_R \\ [\gamma P_S + 1 - \gamma]\dfrac{2}{\theta_R}e^{-\frac{2}{\theta_R}(t_{bc}-\tau_R)}, & t_{bc} > \tau_R \end{cases} \tag{5}$$

$$f(t_{bc}) = \frac{2}{\theta_R}e^{-\frac{2}{\theta_R}(t_{ac}-\tau_R)}, \quad t_{ac} > \tau_R$$

In the case of outflow shown in Figure 1c, the density distribution of coalescent times of each sequence pair is:

$$f(t_{ab}) = \begin{cases} \dfrac{2}{\theta_T} e^{-\frac{2}{\theta_T}(t_{ab}-\tau_T)}, & \tau_T < t_{ab} < \tau_R \\ P_T \dfrac{2}{\theta_R} e^{-\frac{2}{\theta_R}(t_{ab}-\tau_R)}, & t_{ab} > \tau_R \end{cases}$$

$$f(t_{bc}) = \begin{cases} \gamma \dfrac{2}{\theta_S} e^{-\frac{2}{\theta_S}(t_{bc}-\tau_S)}, & \tau_S < t_{bc} < \tau_T \\ \gamma P_S \dfrac{2}{\theta_T} e^{-\frac{2}{\theta_T}(t_{bc}-\tau_T)}, & \tau_T < t_{bc} < \tau_R \\ [\gamma P_S P_T + (1-\gamma)] \dfrac{2}{\theta_R} e^{-\frac{2}{\theta_R}(t_{bc}-\tau_R)}, & t_{bc} > \tau_R \end{cases} \quad (6)$$

$$f(t_{ac}) = \begin{cases} \gamma \dfrac{2}{\theta_T} e^{-\frac{2}{\theta_T}(t_{ac}-\tau_T)}, & \tau_T < t_{ac} < \tau_R \\ [\gamma P_T + (1-\gamma)] \dfrac{2}{\theta_R} e^{-\frac{2}{\theta_R}(t_{ac}-\tau_R)}, & t_{ac} > \tau_R \end{cases}$$

### Supplementary Note 3

We reanalyzed the transcriptome data of *Thuja.* Li et al. (2022) previously estimated the species phylogeny among five *Thuja* species using ASTRAL (Zhang et al. 2018) and used PhyloNet/MPL (Yu and Nakhleh 2015) to identify a ghost introgression event from an unknown ancestral lineage of *Thuja* to *Thuja sutchuenensis*. Here, with an outgroup species *Thujopsis dolabrata*, we focused on three ingroup *Thuja* species, which are *Thuja sutchuenensis* (Tsu), the recipient of the ghost introgression, its sister species *Thuja standishii* (Tst), and a basal species *Thuja plicata* (Tpl) that did not participate in gene flow with the first two species. The one-individual dataset consists of 2969 single-copy non-recombination genes (Li et al. 2022).

Using the backbone topology ((Tsu, Tst), Tpl) from Li et al. (2022), we examined the behaviors of HyDe and PhyloNet/MPL. The jackknife method was used to calculate the P-value for HyDe. For PhyloNet/MPL, we estimated maximum likelihood (ML) gene trees from 2969 one-to-one ortholog alignments of the focal species trio and the outgroup *Solanum lycopersicum* using IQ-TREE v2 (Minh et al. 2020) with ModelFinder (Kalyaanamoorthy et al. 2017) to select the best-fitting substitution model and 1000 ultrafast bootstraps (-B 1000 -m MFP). These unrooted gene trees were then rooted with the outgroup *Solanum lycopersicum* and the outgroup was removed for PhyloNet/MPL. Furthermore, we compared three models of ghost introgression, inflow where gene flow occurs from Tpl to Tst, and outflow in the opposite direcion, using the marginal likelihood values calculated by BPP v4.6.2 (Rannala and Yang 2017; Flouri et al. 2020) with 16 quadrature points. For each of the 16 MCMC runs, we used 50,000 iterations for the burnin, after which we took 1,000,000 posterior samples, sampling every 2 iterations. We then conducted the A00 analysis to estimate parameters for the optimal introgression model. Each run took ~50 hrs using single thread.

The results are presented in Table S4. Firstly, HyDe consistently detected significant gene flow in *Thuja* and inferred the inflow scenario with *Thuja standishii* as the hybrid lineage. In contrast, PhyloNet/MPL failed to distinguish between the three scenarios of ghost introgression, inflow, and outflow as their log pseudo-likelihood values were close to each other. Lastly, we employed BPP to compare the three introgression models. The ghost introgression model was favored with a significant difference in log marginal likelihood value compared to the other models. Our result aligns with the conclusion of the original study by Li et al. (2022).

### Supplementary Figures

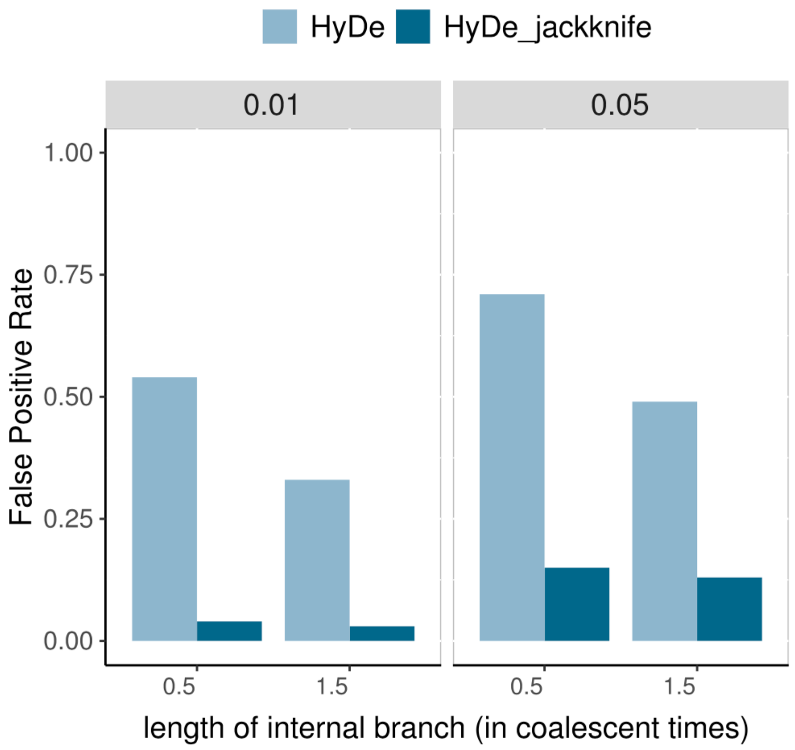


FIGURE S1. High false positive rate of HyDe. For the simulated scenarios shown in Figure 3 with $\gamma = 0$, x-axis represents the internal branch length (in coalescent times) within the speciation history, y-axis represents false positive rate of HyDe and HyDe_jackknife at the significant levels $\alpha = 0.01, 0.05$ among 100 repeats. HyDe_jackknife refers to using block-jackknife procedure for pvalue calculation.

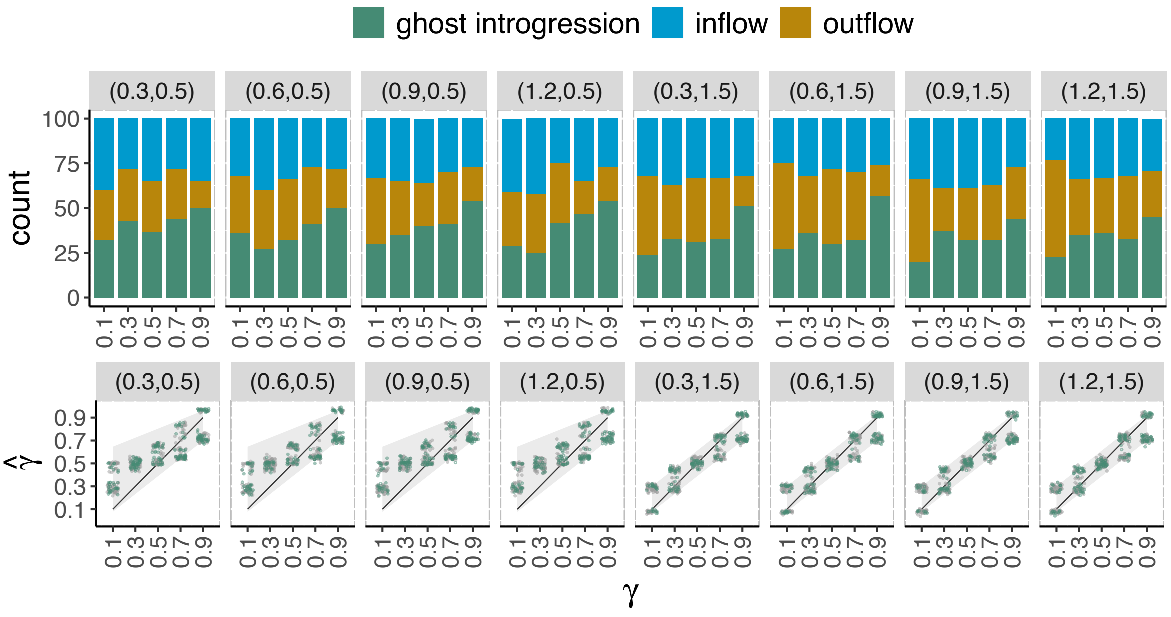


Figure S2. Results of PhyloNet/MPL for 4 sequences per species in the scenarios of ghost introgression. The simulated ghost introgression scenario and corresponding parameter settings are illustrated in Figure 3a. The strips located on the top and right of each plot represent branch lengths in coalescent units $(C_1, C_2)$ and the methods employed, respectively. The x-axis indicates the value of $\gamma$. The figure above represents estimation of network topologies. The numbers of topologies of three network models (i.e., ghost introgression, inflow and outflow) inferred PhyloNet/MPL (using the command InferNetwork_MPL) among 100 replicates are represented by colored bars. The figure below represents estimation of introgression probability $\gamma$. Colored and grey points represent the estimates of $\gamma$ in the true and other two false networks, respectively. These points are horizontally jittered to avoid clutter. The solid line shows the true values, and grey shade area represents the expected range of estimation by PhyloNet/MPL according to equation 2 in the main text.

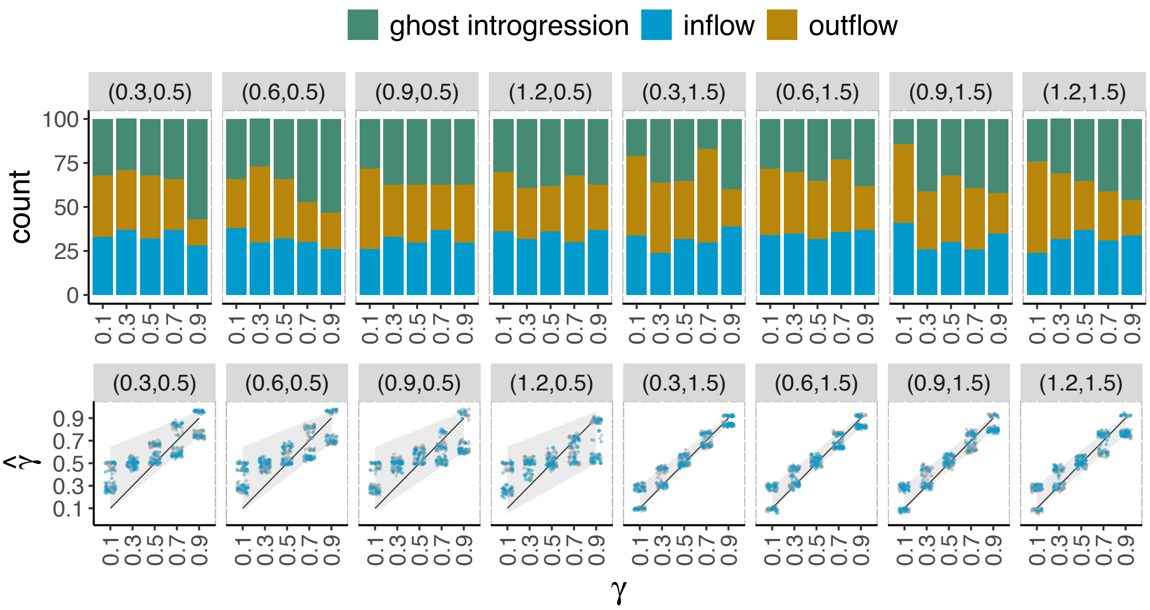


Figure S3. Results of PhyloNet/MPL for 4 sequences per species in the scenarios of inflow introgression. The simulated ghost introgression scenario and corresponding parameter settings are illustrated in Figure 3d. The strips located on the top and right of each plot represent branch lengths in coalescent units $(C_1, C_2)$ and the methods employed, respectively. The x-axis indicates the value of $\gamma$. The figure above represents estimation of network topologies. The numbers of topologies of three network models (i.e., ghost introgression, inflow and outflow) inferred PhyloNet/MPL (using the command InferNetwork_MPL) among 100 replicates are represented by colored bars. The figure below represents estimation of introgression probability $\gamma$. Colored and grey points represent the estimates of $\gamma$ in the true and other two false networks, respectively. These points are horizontally jittered to avoid clutter. The solid line shows the true values, and grey shade area represents the expected range of estimation by PhyloNet/MPL according to equation 2 in the main text.

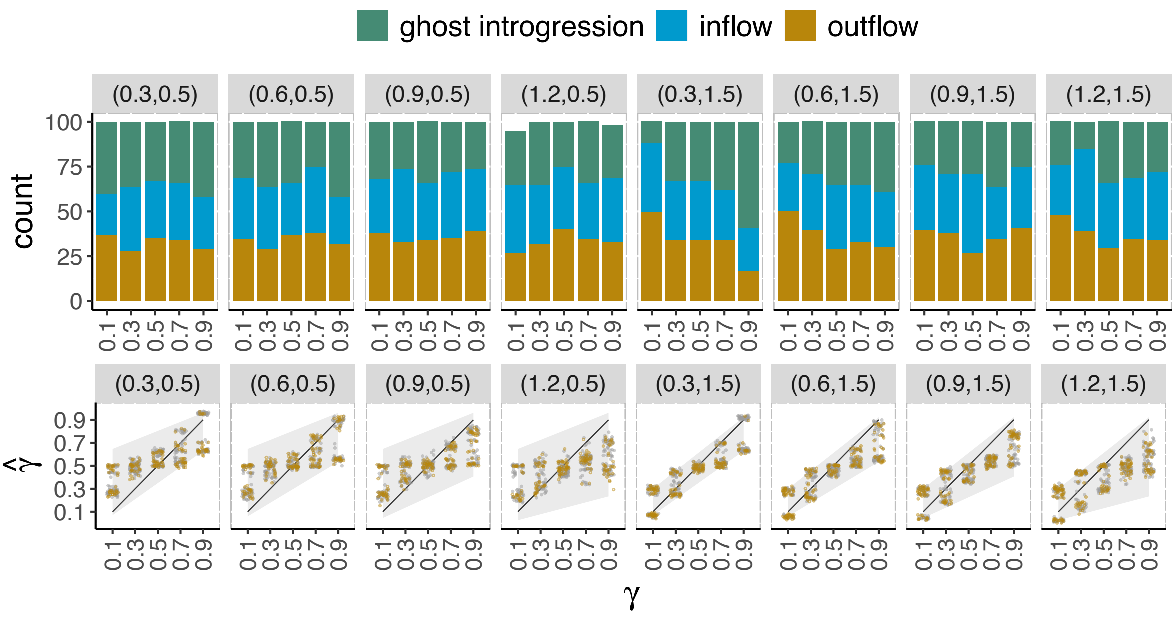


Figure S4. Results of PhyloNet/MPL for 4 sequences per species in the scenarios of outflow introgression. The simulated ghost introgression scenario and corresponding parameter settings are illustrated in Figure 3g. The strips located on the top and right of each plot represent branch lengths in coalescent units $(C_1, C_2)$ and the methods employed, respectively. The x-axis indicates the value of $\gamma$. The figure above represents estimation of network topologies. The numbers of topologies of three network models (i.e., ghost introgression, inflow and outflow) inferred PhyloNet/MPL (using the command InferNetwork_MPL) among 100 replicates are represented by colored bars. The remaining replicates that are not shown correspond to other networks inferred by PhyloNet/MPL. The figure below represents estimation of introgression probability $\gamma$. Colored and grey points represent the estimates of $\gamma$ in the true and other two false networks, respectively. These points are horizontally jittered to avoid clutter. The solid line shows the true values, and grey shade area represents the expected range of estimation by PhyloNet/MPL according to equation 2 in the main text.

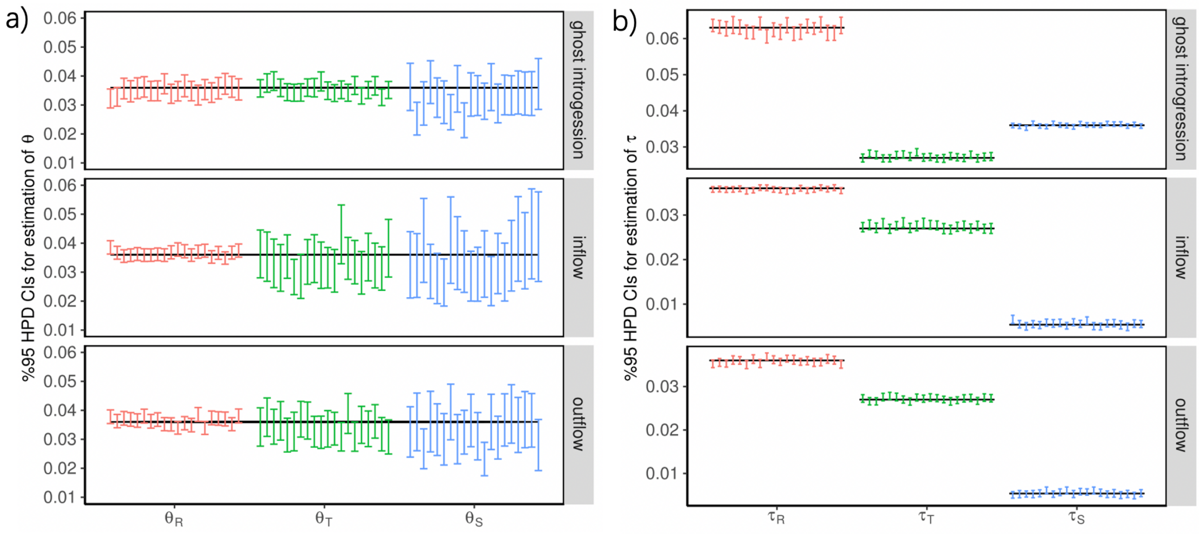


Figure S5. The 95% highest- probability-density (HPD) credibility intervals (CIs) for the parameter $\theta s$ and $\tau s$ by BPP. The scenarios on the strips are illustrated in Figure 3, and are simulated under the parameter combination $(C_1, C_2, \gamma) = (0.3, 0.5, 0.3)$. Black dashed line indicates the true value.

### Supplementary Tables

Table S1. HyDe results for trios that are related to the three introgression events illustrated in Figure 6a

| Introgression | Triple | P1-H-P2 | P-value | Gamma (P1) |
|---|---|---|---|---|
| ① | (Jpro,Jaur,Jcal) | Jpro-Jaur-Jcal | 0.0000000000 | 0.05257135 |
| | (Jpro,Jaur,Jden) | Jpro-Jaur-Jden | 0.0000110667 | 0.02711841 |
| | (Jpro,Jaur,Jgra) | Jpro-Jaur-Jgra | 0.0000001862 | 0.03453010 |
| | (Jpro,Jaur,Jaij) | Jpro-Jaur-Jaij | 0.0000000025 | 0.0432399 |
| | (Jpro,Jaur,Jbif) | Jpro-Jaur-Jbif | 0.0000000006 | 0.0331669 |
| | (Jpro,Jaur,Jinc) | Jpro-Jaur-Jinc | 0.0000000260 | 0.0344734 |
| | (Jpro,Jaur,Jsin) | Jpro-Jaur-Jsin | 0.0000008993 | 0.0305161 |
| | (Jpro,Jaur,Jumb) | Jpro-Jaur-Jumb | 0.0000000000 | 0.0513323 |
| | (Jpro,Jaur,Jqui) | Jpro-Jaur-Jqui | 0.0000000000 | 0.05016790 |
| | (Jpro,Jaur,Jyun) | Jpro-Jaur-Jyun | 0.0000010412 | 0.03141762 |
| | (Jpro,Jaur,Jcal) | Jpro-Jaur-Jcal | 0.0000000000 | 0.05257135 |
| | (Jrep,Jaur,Jaij) | Jrep-Jaur-Jaij | 0.0000000008 | 0.04398558 |
| | (Jrep,Jaur,Jcal) | Jrep-Jaur-Jcal | 0.0000000000 | 0.05102503 |
| | (Jrep,Jaur,Jden) | Jrep-Jaur-Jden | 0.0000215355 | 0.02654750 |
| | (Jrep,Jaur,Jgra) | Jrep-Jaur-Jgra | 0.0000001793 | 0.03287346 |
| | (Jrep,Jaur,Jbif) | Jrep-Jaur-Jbif | 0.0000000000 | 0.0336872 |
| | (Jrep,Jaur,Jinc) | Jrep-Jaur-Jinc | 0.0000000219 | 0.0345052 |
| | (Jrep,Jaur,Jqui) | Jrep-Jaur-Jqui | 0.0000000000 | 0.0480072 |
| | (Jrep,Jaur,Jsin) | Jrep-Jaur-Jsin | 0.0000006671 | 0.0295912 |
| | (Jrep,Jaur,Jyun) | Jrep-Jaur-Jyun | 0.0000000446 | 0.0316709 |
| | (Jrep,Jaur,Jumb) | Jrep-Jaur-Jumb | 0.0000000000 | 0.04937228 |
| | (Jrep,Jaur,Jaij) | Jrep-Jaur-Jaij | 0.0000000008 | 0.04398558 |
| | (Jdar,Jaur,Jaij) | Jdar-Jaur-Jaij | 0.0000000203 | 0.04039135 |
| | (Jdar,Jaur,Jbif) | Jdar-Jaur-Jbif | 0.0000003247 | 0.0311533 |
| | (Jdar,Jaur,Jcal) | Jdar-Jaur-Jcal | 0.0000000000 | 0.0502174 |
| | (Jdar,Jaur,Jden) | Jdar-Jaur-Jden | 0.0000925565 | 0.0236032 |
| | (Jdar,Jaur,Jqui) | Jdar-Jaur-Jqui | 0.0000000000 | 0.0462897 |
| | (Jdar,Jaur,Jsin) | Jdar-Jaur-Jsin | 0.0000262909 | 0.0274957 |
| | (Jdar,Jaur,Jumb) | Jdar-Jaur-Jumb | 0.0000000001 | 0.0472673 |
| | (Jdar,Jaur,Jgra) | Jdar-Jaur-Jgra | 0.0000031959 | 0.03160196 |
| | (Jdar,Jaur,Jinc) | Jdar-Jaur-Jinc | 0.0000017167 | 0.03035070 |
| | (Jdar,Jaur,Jyun) | Jdar-Jaur-Jyun | 0.0000307950 | 0.02805813 |
| ② | (Jdar,Jrep,Jpro) | Jdar-Jrep-Jpro | 0.0000017809 | 0.26571709 |
| ③ | (Jcal,Jumb,Jaij) | Jcal-Jumb-Jaij | 0.0002309009 | 0.0667224 |
| | (Jcal,Jumb,Jbif) | Jcal-Jumb-Jbif | 0.0008170375 | 0.0892047 |

| | | | | |
|---|---|---|---|---|
| | (Jcal,Jumb,Jsin) | Jcal-Jumb-Jsin | 0.0017215347 | 0.0821918 |
| | (Jcal,Jumb,Jden) | Jcal-Jumb-Jden | 0.0000015300 | 0.186301 |
| | (Jcal,Jumb,Jgra) | Jcal-Jumb-Jgra | 0.0000000014 | 0.205043 |
| | (Jcal,Jumb,Jinc) | Jcal-Jumb-Jinc | 0.0000000000 | 0.19059720 |
| | (Jcal,Jumb,Jyun) | Jcal-Jumb-Jyun | 0.0000000000 | 0.45823171 |
| | (Jqui,Jumb,Jaij) | Jqui-Jumb-Jaij | 0.0061468047 | 0.05439864 |
| | (Jqui,Jumb,Jbif) | Jqui-Jumb-Jbif | 0.0071768675 | 0.06454660 |
| | (Jqui,Jumb,Jsin) | Jqui-Jumb-Jsin | 0.0060437127 | 0.0648123 |
| | (Jqui,Jumb,Jden) | Jqui-Jumb-Jden | 0.0000004374 | 0.18066940 |
| | (Jqui,Jumb,Jgra) | Jqui-Jumb-Jgra | 0.0000000000 | 0.18669190 |
| | (Jqui,Jumb,Jinc) | Jqui-Jumb-Jinc | 0.0000000001 | 0.177195 |
| | (Jqui,Jumb,Jyun) | Jqui-Jumb-Jyun | 0.0000000000 | 0.44969851 |

Table S2. PhyloNet/MPL results for trios that are related to the three introgression events illustrated in Figure 6a.

| **Introgression** | **Triple** | **First three optimum scenarios** | **logarithm of pseudo-likelihood** | **Gamma** |
|---|---|---|---|---|
| ① | (Jpro,Jaur,Jcal) | ghost | -5553.977162 | 0.451247 |
| | | outflow | -5553.978751 | 0.428730 |
| | | inflow | -5554.004745 | 0.481771 |
| | (Jpro,Jaur,Jden) | inflow | -5490.284528 | 0.096210 |
| | | outflow | -5490.284701 | 0.086736 |
| | | ghost | -5490.285351 | 0.169099 |
| | (Jpro,Jaur,Jgra) | inflow | -5569.091897 | 0.102636 |
| | | outflow | -5569.092721 | 0.116401 |
| | | ghost | -5569.097411 | 0.189557 |
| | (Jpro,Jaur,Jaij) | outflow | -5600.437188 | 0.110878 |
| | | inflow | -5600.437293 | 0.486673 |
| | | ghost | -5600.440730 | 0.309017 |
| | (Jpro,Jaur,Jbif) | outflow | -5478.377057 | 0.314372 |
| | | ghost | -5478.377764 | 0.076474 |
| | | inflow | -5478.377843 | 0.438345 |
| | (Jpro,Jaur,Jinc) | inflow | -5490.620448 | 0.309017 |
| | | ghost | -5490.620790 | 0.413746 |
| | | outflow | -5490.637295 | 0.099738 |
| | (Jpro,Jaur,Jsin) | inflow | -5454.949213 | 0.181341 |
| | | ghost | -5454.950780 | 0.093361 |
| | | outflow | -5454.955377 | 0.099514 |
| | (Jpro,Jaur,Jumb) | inflow | -5582.779486 | 0.418959 |
| | | ghost | -5582.787039 | 0.215705 |
| | | outflow | -5582.814514 | 0.484520 |
| | (Jpro,Jaur,Jqui) | ghost | -5530.674978 | 0.098661 |
| | | outflow | -5530.675385 | 0.319764 |
| | | inflow | -5530.675616 | 0.204702 |
| | (Jpro,Jaur,Jyun) | inflow | -5554.027527 | 0.444786 |
| | | outflow | -5554.028537 | 0.429043 |
| | | ghost | -5554.068297 | 0.485006 |
| | (Jpro,Jaur,Jcal) | ghost | -5553.977162 | 0.451247 |
| | | outflow | -5553.978751 | 0.428730 |
| | | inflow | -5554.004745 | 0.481771 |
| | (Jrep,Jaur,Jaij) | outflow | -5552.799764 | 0.209687 |
| | | ghost | -5552.799959 | 0.444204 |

| | | | | |
|---|---|---|---|---|
| | | inflow | -5552.803026 | 0.299757 |
| | (Jrep,Jaur,Jcal) | ghost | -5501.698498 | 0.448559 |
| | | inflow | -5501.700333 | 0.414055 |
| | | outflow | -5501.714951 | 0.099676 |
| | (Jrep,Jaur,Jden) | ghost | -5426.123408 | 0.076498 |
| | | inflow | -5426.124227 | 0.152265 |
| | | outflow | -5426.124449 | 0.412196 |
| | (Jrep,Jaur,Jgra) | outflow | -5502.333035 | 0.175974 |
| | | ghost | -5502.341641 | 0.094157 |
| | | inflow | -5502.396126 | 0.474547 |
| | (Jrep,Jaur,Jbif) | inflow | -5408.348709 | 0.416451 |
| | | outflow | -5408.349359 | 0.083468 |
| | | ghost | -5408.349957 | 0.420004 |
| | (Jrep,Jaur,Jinc) | ghost | -5424.144867 | 0.459731 |
| | | inflow | -5424.145059 | 0.415128 |
| | | outflow | -5424.145122 | 0.460410 |
| | (Jrep,Jaur,Jqui) | inflow | -5484.293168 | 0.085962 |
| | | ghost | -5484.293200 | 0.083506 |
| | | outflow | -5484.293270 | 0.443742 |
| | (Jrep,Jaur,Jsin) | ghost | -5415.148006 | 0.075231 |
| | | outflow | -5415.148084 | 0.440266 |
| | | inflow | -5415.149025 | 0.088264 |
| | (Jrep,Jaur,Jyun) | outflow | -5491.194289 | 0.084474 |
| | | ghost | -5491.194428 | 0.100738 |
| | | inflow | -5491.195070 | 0.093056 |
| | (Jrep,Jaur,Jumb) | inflow | -5522.157598 | 0.428255 |
| | | ghost | -5522.158834 | 0.099183 |
| | | outflow | -5522.169600 | 0.099303 |
| | (Jrep,Jaur,Jaij) | outflow | -5552.799764 | 0.209687 |
| | | ghost | -5552.799959 | 0.444204 |
| | | inflow | -5552.803026 | 0.299757 |
| | (Jdar,Jaur,Jaij) | outflow | -5558.384441 | 0.441461 |
| | | inflow | -5558.384630 | 0.446395 |
| | | ghost | -5558.386755 | 0.316927 |
| | (Jdar,Jaur,Jbif) | inflow | -5430.562530 | 0.434053 |
| | | outflow | -5430.562728 | 0.085118 |
| | | ghost | -5430.563678 | 0.155350 |
| | (Jdar,Jaur,Jcal) | inflow | -5513.275578 | 0.097434 |
| | | outflow | -5513.276963 | 0.097327 |
| | | ghost | -5513.277713 | 0.174529 |
| | (Jdar,Jaur,Jden) | outflow | -5471.953844 | 0.309017 |

| | | | | |
|---|---|---|---|---|
| | | ghost | -5471.953877 | 0.092553 |
| | | inflow | -5472.020506 | 0.476365 |
| | (Jdar,Jaur,Jqui) | inflow | -5520.874196 | 0.097252 |
| | | outflow | -5520.874365 | 0.444701 |
| | | ghost | -5520.874432 | 0.096045 |
| | (Jdar,Jaur,Jsin) | ghost | -5412.948057 | 0.412954 |
| | | inflow | -5412.948091 | 0.093326 |
| | | outflow | -5412.949082 | 0.322501 |
| | (Jdar,Jaur,Jumb) | ghost | -5530.154528 | 0.449692 |
| | | outflow | -5530.154684 | 0.415824 |
| | | inflow | -5530.164634 | 0.101719 |
| | (Jdar,Jaur,Jgra) | outflow | -5521.079892 | 0.093425 |
| | | ghost | -5521.134127 | 0.477603 |
| | | inflow | -5521.134835 | 0.477619 |
| | (Jdar,Jaur,Jinc) | inflow | -5452.963087 | 0.089415 |
| | | outflow | -5452.963126 | 0.437150 |
| | | ghost | -5452.963416 | 0.155102 |
| | (Jdar,Jaur,Jyun) | outflow | -5497.579820 | 0.086701 |
| | | inflow | -5497.580097 | 0.089247 |
| | | ghost | -5497.580274 | 0.076733 |
| ② | (Jrep,Jdar,Jpro) | inflow | -6987.352815 | 0.467787 |
| | | outflow | -6987.353569 | 0.393826 |
| | | ghost | -6987.353797 | 0.739419 |
| ③ | (Jcal,Jumb,Jaij) | outflow | -6451.451204 | 0.193893 |
| | | inflow | -6451.451546 | 0.253458 |
| | | ghost | -6451.452710 | 0.286680 |
| | (Jcal,Jumb,Jbif) | inflow | -6758.547385 | 0.560893 |
| | | ghost | -6758.548531 | 0.621506 |
| | | outflow | -6758.549450 | 0.590931 |
| | (Jcal,Jumb,Jden) | ghost | -6877.212403 | 0.430461 |
| | | inflow | -6877.212557 | 0.434090 |
| | | outflow | -6877.213002 | 0.355934 |
| | (Jcal,Jumb,Jgra) | outflow | -6857.187077 | 0.518271 |
| | | inflow | -6857.187742 | 0.506000 |
| | | ghost | -6857.188799 | 0.309017 |
| | (Jcal,Jumb,Jinc) | inflow | -6864.224230 | 0.361667 |
| | | ghost | -6864.224572 | 0.367553 |
| | | outflow | -6864.225318 | 0.630840 |
| | (Jcal,Jumb,Jsin) | ghost | -6650.997747 | 0.539230 |
| | | outflow | -6650.998383 | 0.270320 |
| | | inflow | -6650.998454 | 0.494000 |

| | | | | |
|---|---|---|---|---|
| | (Jcal,Jumb,Jyun) | outflow | -6961.509963 | 0.715410 |
| | | inflow | -6961.511348 | 0.551944 |
| | | ghost | -6961.511625 | 0.455973 |
| | (Jqui,Jumb,Jaij) | inflow | -6361.797703 | 0.241505 |
| | | outflow | -6361.799573 | 0.500000 |
| | | ghost | -6361.801383 | 0.494000 |
| | (Jqui,Jumb,Jbif) | ghost | -6712.026145 | 0.536974 |
| | | inflow | -6712.027415 | 0.304780 |
| | | outflow | -6712.027451 | 0.265059 |
| | (Jqui,Jumb,Jden) | inflow | -6815.272979 | 0.494000 |
| | | ghost | -6815.273088 | 0.346382 |
| | | outflow | -6815.273587 | 0.479836 |
| | (Jqui,Jumb,Jgra) | inflow | -6811.780667 | 0.489344 |
| | | outflow | -6811.781049 | 0.540941 |
| | | ghost | -6811.781063 | 0.386111 |
| | (Jqui,Jumb,Jinc) | outflow | -6825.674326 | 0.625309 |
| | | ghost | -6825.674694 | 0.608135 |
| | | inflow | -6825.674831 | 0.559447 |
| | (Jqui,Jumb,Jsin) | inflow | -6560.942613 | 0.523981 |
| | | ghost | -6560.942795 | 0.500000 |
| | | outflow | -6560.942795 | 0.500000 |
| | (Jqui,Jumb,Jyun) | ghost | -6935.006360 | 0.474611 |
| | | outflow | -6935.008608 | 0.644961 |
| | | inflow | -6935.011324 | 0.617872 |

Table S3. Parameter estimation in BPP for trios that are related to the three introgression events illustrated in Figure 6a. The nodes S, T and R represent the time for the introgression event, the split time of the sister species, and the root, respectively.

| **Introgression** | **Triple** | **The optimum network** | **Node** | **Time ($\tau$)** | **Gamma** |
|---|---|---|---|---|---|
| ① | (Jdar,Jaur,Jcal) | outflow | S | 0.000833 | 0.058379 |
| | | | T | 0.001452 | |
| | | | R | 0.003614 | |
| | (Jdar,Jaur,Jyun) | outflow | S | 0.000831 | 0.050015 |
| | | | T | 0.001364 | |
| | | | R | 0.003461 | |
| ② | (Jrep,Jdar,Jpro) | outflow | S | 0.001715 | 0.110150 |
| | | | T | 0.002109 | |
| | | | R | 0.003082 | |
| ③ | (Jcal,Jumb,Jaij) | outflow | S | 0.000721 | 0.081366 |
| | | | T | 0.001032 | |
| | | | R | 0.001953 | |

Table S4: Detection of ghost introgression for three species *Thuja sutchuenensis* (Tsu), *Thuja standishii* (Tst) and *Thuja plicata* (Tpl)

| **Method** | **Ghost Introgression** 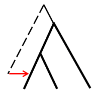  | **Inflow** 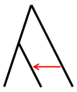  | **Outflow** 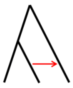  |
|---|---|---|---|
| **HyDe** | | ✓ $\gamma$=0.42 | |
| **PhyloNet/MPL** | ✓ (-3151.965) $\gamma$=0.54 | (-3151.966) | (-3151.966) |
| **BPP** | ✓ (-6159557.47) $\gamma$=0.06 | (-6160280.57) | (-6160083.98) |

Note: the numbers in brackets indicate the logarithm of the pseudo-likelihood values calculated by PhyloNet/MPL and the logarithm of the marginal likelihood calculated by BPP for the three models. $\gamma$ refers to introgression probability.

**References**

Flouri T, Jiao XY, Rannala B, Yang ZH. 2020. A Bayesian implementation of the multispecies coalescent model with introgression for phylogenomic analysis. Mol. Biol. Evol. 37:1211–1223.

Kalyaanamoorthy S, Minh BQ, Wong TK, Von Haeseler A, Jermiin LS. 2017. Modelfinder: Fast model selection for accurate phylogenetic estimates. Nat. Methods 14:587–589.

Li J, Zhang Y, Ruhsam M, Milne RI, Wang Y, Wu D, Jia S, Tao T, Mao K. 2022. Seeing through the hedge: Phylogenomics of *Thuja* (Cupressaceae) reveals prominent incomplete lineage sorting and ancient introgression for Tertiary relict flora. Cladistics 38:187–203.

Minh BQ, Schmidt HA, Chernomor O, Schrempf D, Woodhams MD, von Haeseler A, Lanfear R. 2020. IQ-TREE 2: New models and efficient methods for phylogenetic inference in the genomic era. Mol. Biol. Evol. 37:1530–1534.

Rannala B, Yang Z. 2017. Efficient Bayesian species tree inference under the multispecies coalescent. Syst. Biol. 66:823–842.

Yu Y, Nakhleh L. 2015. A maximum pseudo-likelihood approach for phylogenetic networks. BMC Genom. 16:S10.

Zhang C, Rabiee M, Sayyari E, Mirarab S. 2018. ASTRAL-III: Polynomial time species tree reconstruction from partially resolved gene trees. BMC Bioinform. 19:153.
